# Spatial and Single-Cell Transcriptomics Decipher the Crosstalk Environment of *DEFB1*^+^ Cancer Cells and *IFI30*^+^ Macrophages in Intrahepatic Cholangiocarcinoma

**DOI:** 10.1101/2025.11.22.689892

**Authors:** Guoliang Wang, Hang Meng, Meng Li, Xiangdong Fang, Weilong Zou, Hongzhu Qu

## Abstract

Intrahepatic cholangiocarcinoma (ICC) is a rare but highly aggressive primary liver malignancy distinguished by a profoundly heterogeneous tumor microenvironment, which underlies its limited response to targeted and immune-based therapies. Deciphering the cellular composition and intercellular signaling within this complex ecosystem is critical to understanding ICC progression and to identifying actionable therapeutic targets. Here, we integrated single-cell RNA sequencing and spatial transcriptomics to construct a comprehensive cellular and transcriptional atlas of human ICC. Our analyses revealed a continuous trajectory of T-cell state transition from activation to exhaustion, with *CD8*^+^ proliferating T cells exhibiting two distinct exhaustion programs, namely terminal and progenitor-like exhaustion. Notably, *DEFB1*^+^ cholangiocytes and *IFI30*^+^ macrophages displayed a strong positive correlation across independent ICC cohorts and were found in close spatial proximity within the tumor microenvironment. Mechanistically, their interaction appears to be mediated by the *TGFB1* – *TGFBR1* signaling pathway, contributing to tumor progression and immunotherapy resistance. Together, these findings delineate the cellular architecture and spatial organization of the ICC microenvironment, uncovering a key cholangiocyte–macrophage axis that shapes immune dysfunction and offers a potential therapeutic entry point to enhance immunotherapy efficacy in ICC.

## Introduction

As the second most common primary hepatic malignancy, intrahepatic cholangiocarcinoma (ICC) constitutes approximately 20% of all liver cancer cases [1]. Characterized by high aggressiveness and inherent chemoresistance, this malignancy has witnessed a consistent global upward trend in incidence over the past decade, with its prognostic outlook remaining dismal [2]. Radical surgical resection persists as the only potential curative modality for ICC. Nevertheless, the 5-year survival rate postoperatively remains merely around 20% [3]. Furthermore, the majority of patients present with advanced-stage at the time of diagnosis, at which point the optimal therapeutic window for radical resection has already elapsed [4, 5].

In recent years, immunotherapy has revolutionized the landscape of cancer therapeutics. Immune checkpoint blockade (ICB) employs monoclonal antibodies to targets inhibitory receptors, including programmed cell death 1 (PD-1), programmed cell death ligand 1 (PD-L1), and cytotoxic T-lymphocyte antigen 4 (CTLA-4), with the primary goal of reinstating the antitumor effector function of T cells. Notably, ICB has demonstrated remarkable clinical efficacy in a spectrum of refractory malignancies, such as advanced hepatocellular carcinoma [6, 7]. In contrast, the majority of ICC patients attain only modest clinical benefits from these regimens. This discrepancy underscores the imperative to dissect the cellular and spatial heterogeneity of ICC, as well as the intricate crosstalk between ICC cells and the immune microenvironment [8, 9]. Such mechanistic insights may uncover novel and more effective therapeutic targets, thereby optimizing the clinical outcomes of immunotherapy in ICC.

Single-cell RNA sequencing (scRNA-seq) facilitates high-resolution profiling of intra- and intercellular molecular dynamics, which has been extensively employed to decipher cellular heterogeneity within and between tumors [10, 11]. Although single-cell RNA sequencing (scRNA-seq) provides unprecedented resolution in dissecting cellular heterogeneity, tissue dissociation inevitably eliminates spatial context, preventing accurate characterization of the spatial organization and interactions among tumor, immune, and stromal cells within the tumor microenvironment (TME)[12, 13]. Spatial transcriptomics (ST), by contrast, enables the concurrent measurement of gene expression and the precise mapping of cellular localization within the TME. This approach is critical for understanding tumor–stromal crosstalk [14–16]. Consequently, the integration of scRNA-seq with ST holds substantial promise for deconvoluting the intercellular communication networks that orchestrate the pathophysiology of ICC. Recent investigations leveraging spatial omics technologies have shed light on the crosstalk between ICC cells and tumor-associated macrophages (TAMs), demonstrating that elevated *TFF3* expression in ICC cells may promote M2-like TAM polarization [17]. Nevertheless, the heterogeneity of both tumor cells and TAMs, as well as the underpinning molecular mechanisms governing their reciprocal interactions, remain incompletely elucidated.

In this study, we integrated droplet-based scRNA-seq with the Visium spatial transcriptomics platform to systematically profile primary ICC tumors and their matched paracancerous tissues. Our comprehensive analyses revealed a highly heterogeneous tumor microenvironment within ICC lesions. We identified two distinct exhaustion patterns among proliferating *CD8*^+^ T cells, namely terminal exhaustion and progenitor cell exhaustion. We also observed a strong positive correlation between *DEFB1*^+^ cholangiocytes and *IFI30*^+^ macrophages, and this correlation was significantly associated with poor overall survival in ICC patients. Spatial transcriptomics analyses confirmed the close colocalization of these two cell populations within the TME. Additionally, cell-cell interaction analysis indicated enrichment of the *TGFB1*–*TGFBR1* signaling axis in their interaction. Collectively, our findings reveal the intricate intercellular interactions between cholangiocytes subsets and macrophage subpopulations, and highlight potential therapeutic targets that may improve the efficacy of immunotherapy for this refractory malignancy.

## Results

### Single-cell transcriptome heterogeneity in ICC

To explore cellular diversity and transcriptomic heterogeneity in ICC, we collected tumor tissue, tumor-adjacent tissue, and distal normal tissue from a patient. Single-cell RNA sequencing (scRNA-seq) was performed using the 10x Genomics platform. In parallel, we integrated 16 scRNA-seq datasets from public databases (Figure 1A). Following rigorous quality control, 119,618 cells were retained for downstream analyses. Dimensionality reduction and clustering identified 29 distinct clusters. Expression patterns of *PTPRC*, *MGP*, and *EPCAM* delineated the major immune, stromal, and epithelial populations in ICC tissues (Figure 1C). Leveraging marker gene signatures, we performed a comprehensive annotation that resolved these populations into 14 major cell types. These major cell types include T cells (50,365 cells; *CD3D^+^*and *CD3E^+^*), bile duct cells (25,837 cells; *EPCAM^+^*, *KRT19^+^*, and *KRT7^+^*), endothelial cells (10,841 cells; *PTPRB^+^*, *CLDN5^+^*, and *CDH5^+^*), monocytes (6,843 cells; *FCN1^+^*, *S100A8^+^*, and *S100A9^+^*), NK cells (5,854 cells; *NCAM1^+^*), macrophages (5,243 cells; *C1QA^+^* and *C1QB^+^*), fibroblasts (4,998 cells; *ACTA2^+^* and *COL1A2^+^*), B cells (2,847 cells; *MS4A1^+^* and *CD79A^+^*), cDC2 (2,177 cells; *CD1A^+^*, *CD1C^+^*, and *CD1E^+^*), hepatocytes (1,550 cells; *APOA1^+^*, *APOC3^+^*, and *FABP1^+^*), plasma cells (1,183 cells; *MZB1^+^*, *IGKC^+^*, and *JCHAIN^+^*), cDC1 (1,122 cells; *CLEC9A^+^*, *XCR1^+^*, and *BATF3^+^*), pDC (493 cells; *LILRA4^+^* and *CLEC4C^+^*), and mast cells (265 cells; *TPSAB1^+^* and *TPSB2^+^*) (Figure 1B). The accuracy of cell identity was confirmed through the expression of marker genes (Figure 1D) and the differentially expressed genes (DEGs) (Figure 1E). Fourteen major cell types were present in tumors and adjacent normal tissue in all patients, although the extent of infiltration of each major cell type varied (Figure 1F). We observed a significant decrease of lymphocytes (T cells, NK cells, and B cells) in ICC, while stromal cells (fibroblasts and endothelial cells) and epithelial cells (cholangiocytes and hepatocytes) showed the increasing trend (Figure 1G, H). These observations suggest that immune infiltration into ICC tumors may be suppressed by the tumor microenvironment.

**Figure 1.**
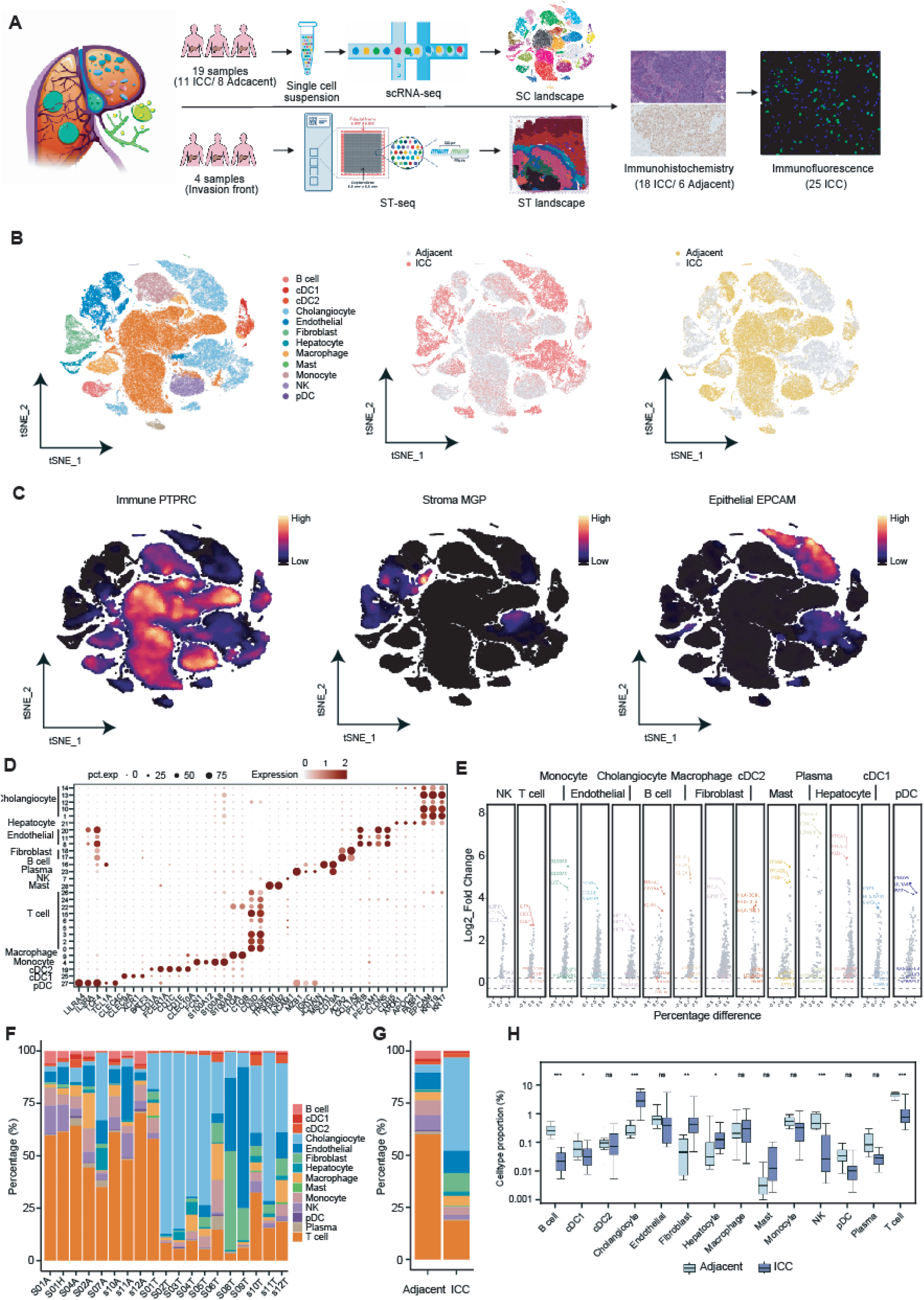
Single-cell transcriptome heterogeneity in ICC. **A.** Graphical overview of the study design. adjacent and tumor tissues from ICC patients were processed into single-cell suspensions, and unsorted cells were used for scRNA-seq by 10x Genomics. Tumor slides were processed using 10x Genomics Visium to obtain spatial transcriptomics. The results were validated using immunohistochemistry and multiplex immunofluorescence. **B.** tSNE plots showing cell types of 119,618 high-quality single cells, split by tissue type. The left panel shows the 14 cell types with colored-coded. The middle and right panels show cells from tumor and adjacent tissues. **C.** tSNE plots showing the expression of known markers for immune cells, stromal cells, and epithelial cells. *PTPRC*, *MGP*, and *EPCAM* represent marker gene highly expressed in immune cells, stromal cells, and epithelial cells, respectively. **D.** The average expression of known markers in cell clusters. The numbers in y-axis mean the 29 cell clusters initially identified. The size of the dots represents the percentage of cells expressing the gene within each cluster. **E.** Volcano plots showing the top differentially expressed genes (DEGs) in each cell type. **F.** The proportion of cell types in each sample. **G.** The proportion of cell types in adjacent and tumor tissues. Cells are colored identically to those shown in panel F. **H.** The differences in the proportions of different cell types between adjacent and tumor tissues. *P* values were determined by T-test. *, *P* < 0.05. **, *P* < 0.01. ***, *P* < 0.001.

### Spatial transcriptome heterogeneity of ICC

To systematically characterize the complex spatial transcriptome structure of intrahepatic cholangiocarcinoma, we collected tissue section from 1 patient (Sample ICC1), generated spatial expression profiles using 10x Visium technology, and integrated spatial transcriptome samples from 3 cases in public databases (Samples ICC2–4). Initially, board-certified anatomical pathologists characterized a total of five distinct histological regions, including tumor region (T), tumor-peritumor junctional region (J), paratumor region (P), tumor stromal region (TS), and paratumor stromal region (PS), within the sections sourced from four patients. Sample ICC1 and ICC3 exhibited the presence of all five histological regions (Figure 2A and Supplementary Figure 1A), while sample ICC2 was devoid of the normal stromal region and sample ICC4 lacked the tumor stromal region (Supplementary Figure1A). To further validate these annotations, we calculated ESTIMATE scores, including immune, stromal, and tumor purity scores. Across the four samples, stromal scores highlighted extensive fibrosis, and tumor purity scores accurately delineated tumor areas. Immune scores were generally low, particularly within tumor regions (Figure 2B, Supplementary Figure 1B). GSVA enrichment analysis of the integrated spatial data revealed that tumor regions were enriched in proliferation-related pathways (G2M checkpoint, E2F targets) and glycolysis (Figure 2C). Glycolysis scoring confirmed higher activity in tumor regions across all four patients, with key glycolytic genes *PFKM* and *PKM* highly expressed (Figure 2D, Supplementary Figure 2). These results suggest that enhanced glycolytic activity in tumor areas may support the energy demands of cell proliferation and immune evasion.

**Figure 2.**
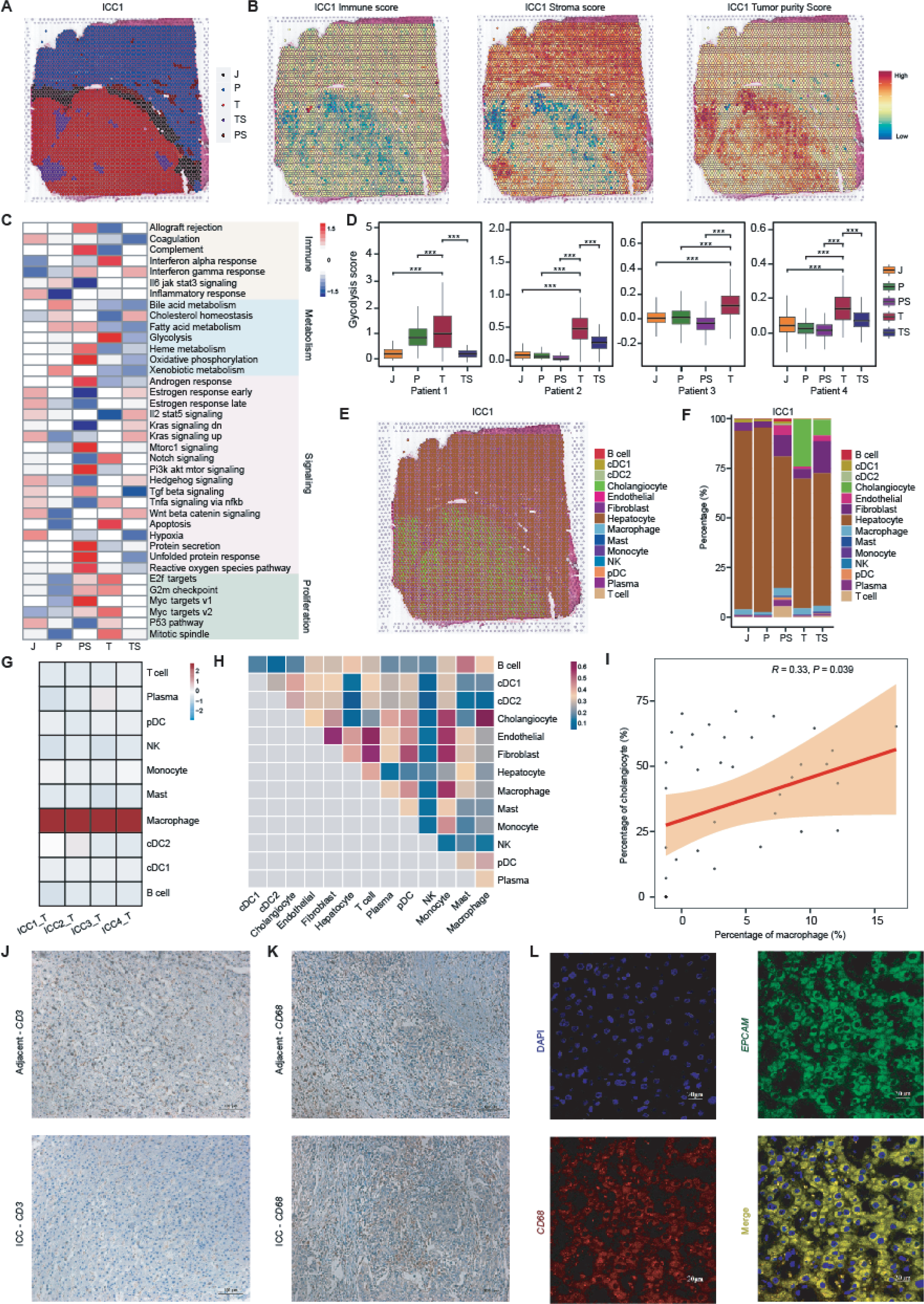
Spatial transcriptome heterogeneity of ICC. **A.** Pathologists manually annotated the tumor area, adjacent area, invasion area, tumor stroma, and adjacent stroma in the paraffin section of patient 1. J, tumor-peritumor junctional region. P, paratumor region. T, tumor region. TS, tumor stroma region. PS, paratumor stroma region. **B.** Expression quantification for each spatial feature was assessed to generate stromal score, immune score, and tumor purity score using ESTIMATE. **C.** Differences in pathway activity (scored per cell by GSVA) across five tissue regions. Blue means low activity and red represents high activity. **D.** The differences in glycolysis levels across different tissue regions in four spatial ICC samples. *P* values were determined by T-test. ***, *P* < 0.001. **E.** The spatial distribution of cells within the ICC1 sample deconvolved using the cell markers identified in single-cell RNA-seq data. **F.** The proportion of cell types in different tissue regions in the spatial sample ICC1. **G.** Heat map showing the degree of immune cell infiltration in tumor region in the four spatial ICC samples. **H.** The spatial co-localization of paired cell types in the spatial sample ICC1. **I.** Correlation of macrophage and cholangiocyte infiltration scores in the TCGA CHOL cohort (n = 50). **J.** The expression of *CD3* (T cell marker) in ICC tissue and adjacent tissue by the immunohistochemical examination. **K.** The expression of *CD68* (macrophages marker) in ICC tissue and adjacent tissue by the immunohistochemical examination. **L.** Single channel image for multiplexed immunofluorescence staining of *EPCAM* and *CD68*.

We performed deconvolution analysis with the integrated single-cell transcriptome data to determine the cellular proportions of spatial transcriptome data. As anticipated, spots in the T zone exhibited a higher proportion of single cell type, specifically cholangiocytes, while spots in the other zones contained a mixture of different cell types, including stromal cells and hepatocytes (Figure 2E, Supplementary Figure 3A). Notably, lymphocytes (T cells and B cells, etc.) were predominantly concentrated in non-tumor areas but significantly diminished in tumor areas, indicating that ICC exhibits characteristics of a cold TME as analyzed through spatial omics (Figure 2F, Supplementary Figure 3B). This observation was confirmed by immunohistochemistry (IHC), which showed higher CD3 expression in adjacent tissue compared to tumor tissue across three independent ICC sections (Figure 2J, Supplementary Figure 3C, Supplementary Figure 4A). Macrophages were the only immune cells infiltrating tumor regions at high levels, suggesting a key role in forming an immunosuppressive microenvironment (Figure 2G). IHC staining confirmed that high CD68 expression in tumor and adjacent tissues from three additional sections (Figure 2K, Supplementary Figure 3D, Supplementary Figure 4A). And the IHC staining of M2 macrophage marker CD163 further verified that tumor-infiltrating macrophages were immunosuppressive (Supplementary Figure 4). Spatial colocalization analysis revealed consistent proximity between cholangiocytes and macrophages (Figure 2H, Supplementary Figure 3E). Pairwise Pearson correlation analysis of the 14 major cell types in ICC tumors from the TCGA CHOL cohort, using CIBERSORTx, showed a significant positive correlation between cholangiocytes and macrophages (Pearson correlation coefficient = 0.33, *P* = 0.039) (Figure 2I). Immunofluorescence labeling confirmed the close spatial association of cholangiocytes (*EPCAM*^+^) and macrophages (*CD68*^+^) in ICC tissues (Figure 2L, Supplementary Figure 3F). Together, these findings suggest potential cellular interactions between cholangiocytes and macrophages, which may play a significant role in driving ICC progression.

### Heterogeneous exhaustion patterns of *CD8*^+^ Tpr populations in ICC patients

T lymphocytes within tumor tissues exhibit a high degree of heterogeneity and play a pivotal role in immune evasion and response to immunotherapy [18, 19]. In this study, we observed eight distinct subclusters of T lymphocytes: *CD4*^+^ naive T cells (*CD4*^+^ Tn), *CD4*^+^ effector memory T cells (*CD4*^+^ Tem), regulatory T cells (Treg), *CD8*^+^ naive T cells (*CD8*^+^ Tn), *CD8*^+^ tissue-resident T cells (*CD8*^+^ Trm), *CD8*^+^ effector memory T cells (*CD8*^+^ Tem), *CD8*^+^ terminally differentiated effector memory T cells (*CD8*^+^ Temra), and *CD8*^+^ proliferating T cells (*CD8*^+^ Tpr) based on the expression of marker genes (Figure 3A, Supplementary Figure 5A). In tumor tissues, there was a significant increase in Treg cell numbers (Figure 3B, C, Supplementary Figure 5B), characterized by pronounced expression of immunosuppressive markers such as *TIGIT*, *TNFRSF4*, *CTLA4*, *IL2RA*, and *TNFRSF18* (Supplementary Figure 5A). Additionally, we observed an increasing trend in *CD8*^+^ proliferating T cells (Figure 3B, C, Supplementary Figure 5B), which expressed a number of exhaustion markers, including *LAG3*, *TIGIT*, *HAVCR2*, and *SIRPG*, indicating an exhaustion state for these cells (Figure 3D). Trajectory analysis of *CD4*^+^ T cells revealed that naive T cells occupied the early stage and progressively differentiated into effector memory and regulatory T cells (Figure 3E). The immunosuppressive molecules *TNFRSF4*, *IL2RA*, and *TNFRSF18* were predominantly expressed at the terminal stage of differentiation (Supplementary Figure 5C). *CD8*^+^ T cells displayed a similar pseudo-time progression, beginning with naive and tissue-resident cells, transitioning through effector memory cells, and culminating in terminally differentiated and proliferating subsets. This pattern reflects a state transition from activation to exhaustion (Figure 3F).

**Figure 3.**
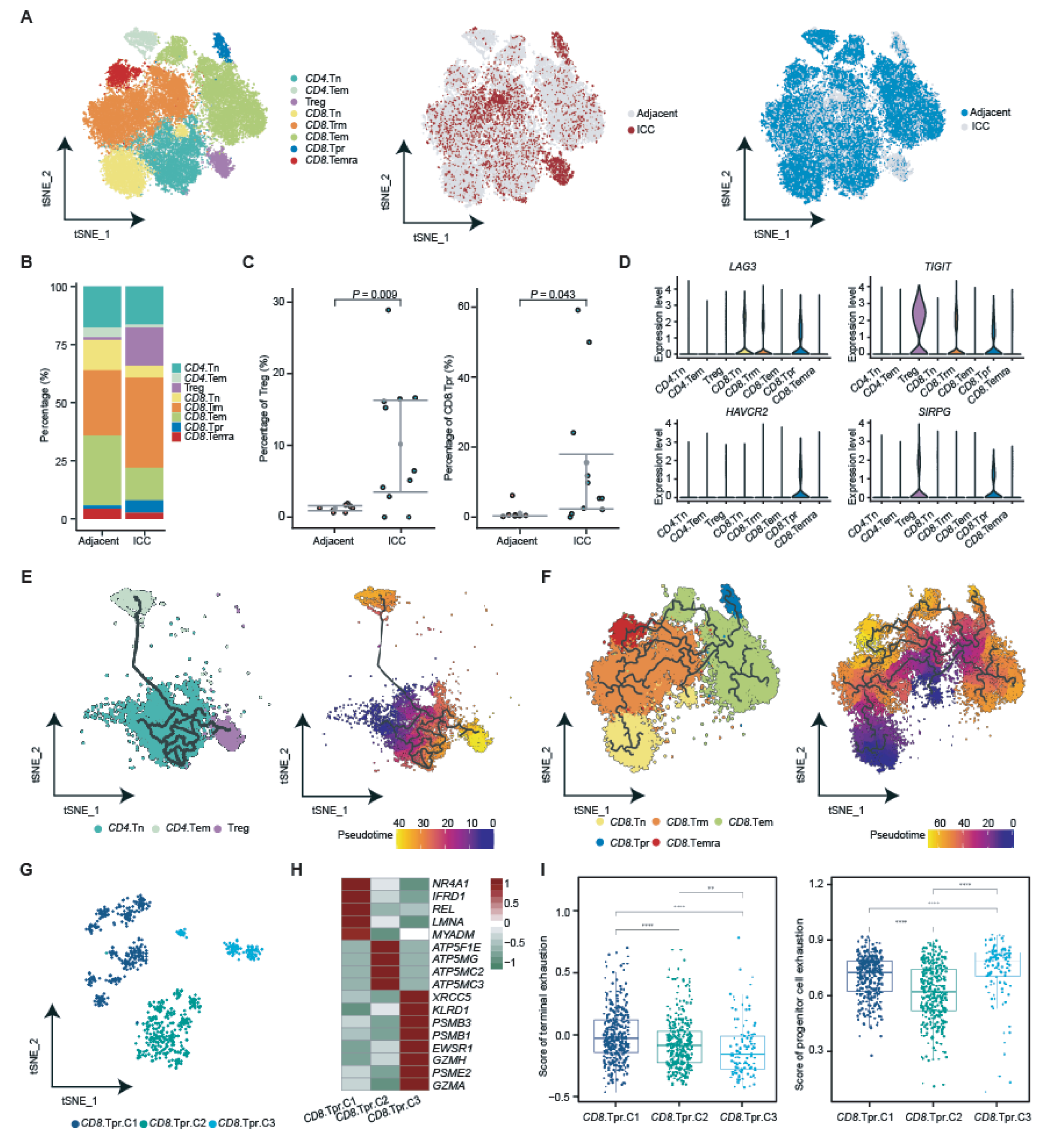
Different exhaustion patterns of *CD8*^+^ Tpr subclusters in ICC patients. **A.** The distribution of T cell subclusters. The left tSNE plot showing the composition of T cells, colored by cluster. The middle and right panels show cells from tumor and adjacent tissues. **B.** The proportion of T cell subtypes in adjacent and tumor tissues. **C.** The difference in the proportion of regulatory T cells (left) and proliferating *CD8*^+^ T cells (right) between adjacent and tumor tissues. *P* values were determined by T-test. **D.** The expression of exhaustion-related genes in different T cell subclusters. **E.** tSNE plot showing the differentiation trajectory of *CD4*^+^ T cells. The left plot shows the trajectory among different types of cells and the right plot shows the trajectory. **F.** tSNE plot showing the differentiation trajectory of *CD8*^+^ T cells. The left plot shows the trajectory among different types of cells and the right plot shows the trajectory. **G.** The composition of proliferating *CD8*^+^ T cells, colored by subclusters. **H.** The expression levels of top DEGs in different proliferating *CD8*^+^ T cell clusters. **I.** The terminal exhaustion score (left) and progenitor cell exhaustion score (right) in different proliferating *CD8*^+^ T cell clusters. *P* values were determined by T-test. *, *P* < 0.05. **, *P* < 0.01. ***, *P* < 0.001. ****, *P* < 0.0001.

Given the spatial transcriptome deconvolution results indicating that *CD8*^+^ proliferating T cells were the most abundant T cell subset infiltrating tumor tissue (Supplementary Figure 5D), we further subdivided *CD8*^+^ Tpr into three clusters (C1-C3) to explore their potential heterogeneity (Figure 3G). *CD8*^+^ Tpr-C1, *CD8*^+^ Tpr-C2, and *CD8*^+^ Tpr-C3 exhibited increased expression of terminal exhaustion marker genes, mitochondrial ATP synthesis genes, and progenitor cell exhaustion genes, respectively (Figure 3H). Previous studies have reported that in cancer immunotherapy models, *CD8*^+^ T cells include a subset containing a progenitor-depleted population that comprises self-renewing cells promoting ICI-induced tumor control and ultimately differentiating into a terminally exhausted population [20]. To verify whether *CD8*^+^ Tpr-C3 exhibits a progenitor cell exhaustion phenotype, we calculated the terminal and progenitor cell exhaustion scores for each *CD8*^+^ cell cluster and found that *CD8*^+^ Tpr-C1 was significantly enriched in the signature of terminal exhaustion. In contrast, *CD8*^+^ Tpr-C3 exhibited significantly higher activity in progenitor cell exhaustion characteristics (Figure 3I). Collectively, these observations suggest two heterogeneous patterns of *CD8*^+^ proliferating T cell exhaustion in ICC, both potentially contributing to ICC tumor progression.

### *DEBF1*^+^ malignant cells are associated with ICC progression

Since ICC originates from cholangiocytes [21], we next examined their composition. The 25,837 cholangiocytes were subdivided into 6 subclusters, including *DEFB1*^+^, *PSCA*^+^, *CCL4*^+^, *PTTG1*^+^, *CCL21*^+^, and *TFFs*^+^ cholangiocytes (Figure 4A, Supplementary Figure 6A), named by the most significant DEGs in subclusters (Supplementary Figure 6B). These subclusters displayed distinct gene expression patterns and biological functions, with proportions varying substantially among patients, reflecting high intra- and inter-tumor heterogeneity (Figure 4B, C). *DEFB1*^+^ cholangiocytes is the predominant subpopulation in ICC, as confirmed by spatial deconvolution analysis (Figure 4D, Supplementary Figure 6D). Immunofluorescence labeling further confirmed the presence of DEFB1^+^ cholangiocytes in tumor tissues (Figure 4M, Supplementary Figure 7). *PSCA*^+^ cholangiocytes almost exclusively exist in tumor tissues and are nearly absent in normal controls, indicating their status as tumor cells with high proliferation and differentiation capabilities (Supplementary Figure 6C), consistent with the fact that they are enriched in epithelial cell differentiation, vascular development, and P53 downstream pathways in PID (Figure 4E). In contrast, *CCL4*^+^ cholangiocytes represent the predominant ductal cell population in normal control tissue, indicating the minimal malignancy of these cells (termed “normal cholangiocytes”) (Supplementary Figure 6C). Compared with paracancerous tissue, *DEFB1*^+^, *PSCA*^+^, and *PTTG1*^+^ cholangiocytes were increased in tumor tissues, while *CCL4*^+^, *CCL21*^+^, and *TFFs*^+^ cholangiocytes were decreased (Figure 4B, C).

**Figure 4.**
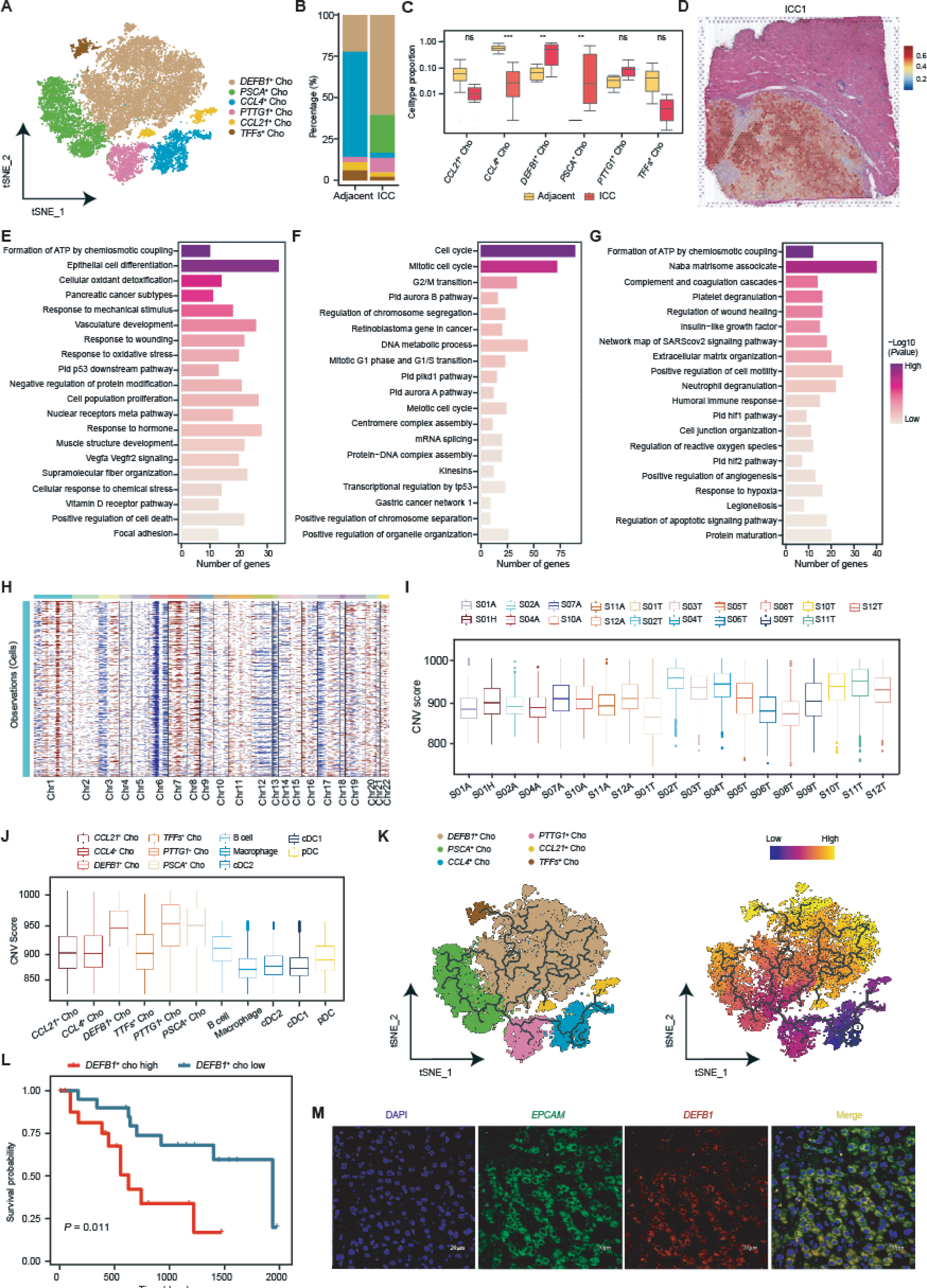
*DEBF1*^+^ malignant cells are associated with ICC progression. **A.** tSNE plot showing cholangiocytes composition, colored by cluster. **B.** The proportion of cholangiocyte subclusters in adjacent and tumor tissues. Cells are colored identically to those shown in panel A. **C.** The differences in the proportions of different cell types between adjacent and tumor tissues. *P* values were determined by T-test. *, *P* < 0.05. **, *P* < 0.01. ***, *P* < 0.001. **D.** The distribution of *DEFB1*^+^ cholangiocytes in the spatial sample ICC1. **E-G.** The enriched KEGG pathways of DEGs in *PSCA*^+^ cholangiocytes (E), *PTTG1*^+^ cholangiocytes (F), and ***DEFB1***^+^ cholangiocytes (G), respectively. **H.** Chromosomal copy number variations of cholangiocytes. **I.** The CNV signals for each sample. **J.** The CNV signals for each cell type. **K.** The differentiation trajectory of cholangiocyte clusters. **L.** Survival analysis of *DEFB1*^+^ cholangiocytes in the TCGA CHOL cohort (n = 50). The mean *DEFB1*^+^ cholangiocytes score provided by coxph is used to split samples. Samples with score greater than the mean score of all samples are labeled as “*DEFB1*^+^ cho high”, and the others are labeled as “*DEFB1*^+^ cho low”. *P* values were determined by T-test. **M.** Single channel image for multiplexed IF staining of EPCAM and DEFB1.

To assess the malignant status of these subpopulations, we inferred single-cell copy number variation (CNV) profiles using inferCNV, with myeloid cells and B cells as references. High CNV accumulation was enriched in certain chromosomes, such as chr7 and chr8 (Figure 4H, I). *DEFB1*^+^, *PSCA*^+^, and *PTTG1*^+^ cholangiocytes exhibited higher CNV scores, along with elevated expression of oncogenes (*KRAS*, *MYC*, *CCND1*, and *EGFR*) and proliferation-related genes (*MKI67* and *TOP2A*), supporting their classification as malignant tumor cells (Figure 4J, Supplementary Figure 6E). Malignant *PTTG1*^+^ cholangiocytes appeared at an earlier stage of cell trajectory, suggesting highly proliferative characteristics (Figure 4K). The DEGs of *PTTG1*^+^ cholangiocytes were enriched in cell cycle, G2/M transition, mitotic G1 phase and G1/S transition, and meiotic cell cycle (Figure 4F). Malignant *DEFB1*^+^ cholangiocytes were positioned in the terminal stage of trajectory differentiation, highly expressing genes involved in multiple pathways related to tumor progression, including regulation of wound healing, positive regulation of angiogenesis, inflammatory response, hypoxia response, and reactive oxygen metabolism process (Figure 4G). Using CIBERSORTx to analyze the TCGA CHOL cohort, we found that *DEFB1*^+^ cholangiocytes were significantly enriched in tumor tissues (Supplementary Figure 6F). Importantly, patients with higher *DEFB1*^+^ cholangiocyte infiltration had shorter overall survival compared with those with lower infiltration, a finding validated in the E-MTAB-6389 cohort (Figure 4L, Supplementary Figure 6G).

### *IFI30*^+^ macrophages are associated with ICC progression

Macrophages were subdivided into seven subtypes: *IFI30*^+^ Mac, *IL8*^+^ Mac, *CD3E*^+^ Mac, *FOLR2*^+^ Mac, *STMN1*^+^ Mac, *KRTs*^+^ Mac, and *CCL14*^+^ Mac (Figure 5A). Among these, *IFI30*^+^ macrophages emerge as the predominant macrophage subpopulation within tumor tissues (Figure 5B), exhibiting high expression of the lysosomal thiol reductase (*IFI30*), steopontin-encoding gene (*SPP1*) and Fibronectin 1 (*FN1*) as illustrated in Supplementary Figure 8A. In contrast, *FOLR2*^+^ macrophages, abundant in normal and adjacent tissues (Figure 5B), represent a type of tissue-resident macophage, expressing non-canonical myeloid markers (*LYVE1* and *SELENOP*) alongside heightened levels of complement genes (*C1QA* and *C1QC*) and scavenger receptors (*MACRO* and *SEPP1*), as depicted in Supplementary Figure 8A. *CD3E*^+^ macrophages highly express immune regulatory molecules (*CD3E* and *CD7*), chemokines (*CCL5* and *IL32*), and calcium-binding proteins (*S100A8*, *S100A9*, and *S100A12*), potentially orchestrating inflammatory and immune response. Within *KRTs*^+^ macrophages, keratin-encoding genes (*KRT8*, *KRT18*, and *KRT19*) are significantly expressed (Supplementary Figure 8A). Previous studies categorize macrophages into M1/M2 polarization states in vitro [22]. We assessed the expression of M1 (Figure 5C), M2 (Figure 5D), angiogenesis (Figure 5E), and phagocytosis (Figure 5F) gene sets across each macrophage subset. Notably, *IFI30*^+^ macrophages displayed heightened expression of M2-like and angiogenic gene sets, suggesting their potential role as tumor-associated macrophages that facilitate tumor growth while suppressing anti-tumor immunity. Conversely, tissue-resident *FOLR2*^+^ macrophages exhibited superior phagocytic function, suggesting their capacity for anti-tumor effects in ICC by engulfing and clearing tumor cells, consistent with prior findings by Julie Helft [23]. Intriguingly, *FOLR2*^+^ macrophages also displayed a more immunosuppressive M2-like gene set. Previous studies have shown that in hepatocellular carcinoma (HCC), *FOLR2*^+^ tumor-associated macrophages exhibit an immunosuppressive phenotype, thereby promoting HCC progression [24]. These findings imply *FOLR2*^+^ macrophages progressive transition towards a pro-tumor phenotype within the ICC tumor microenvironment.

**Figure 5.**
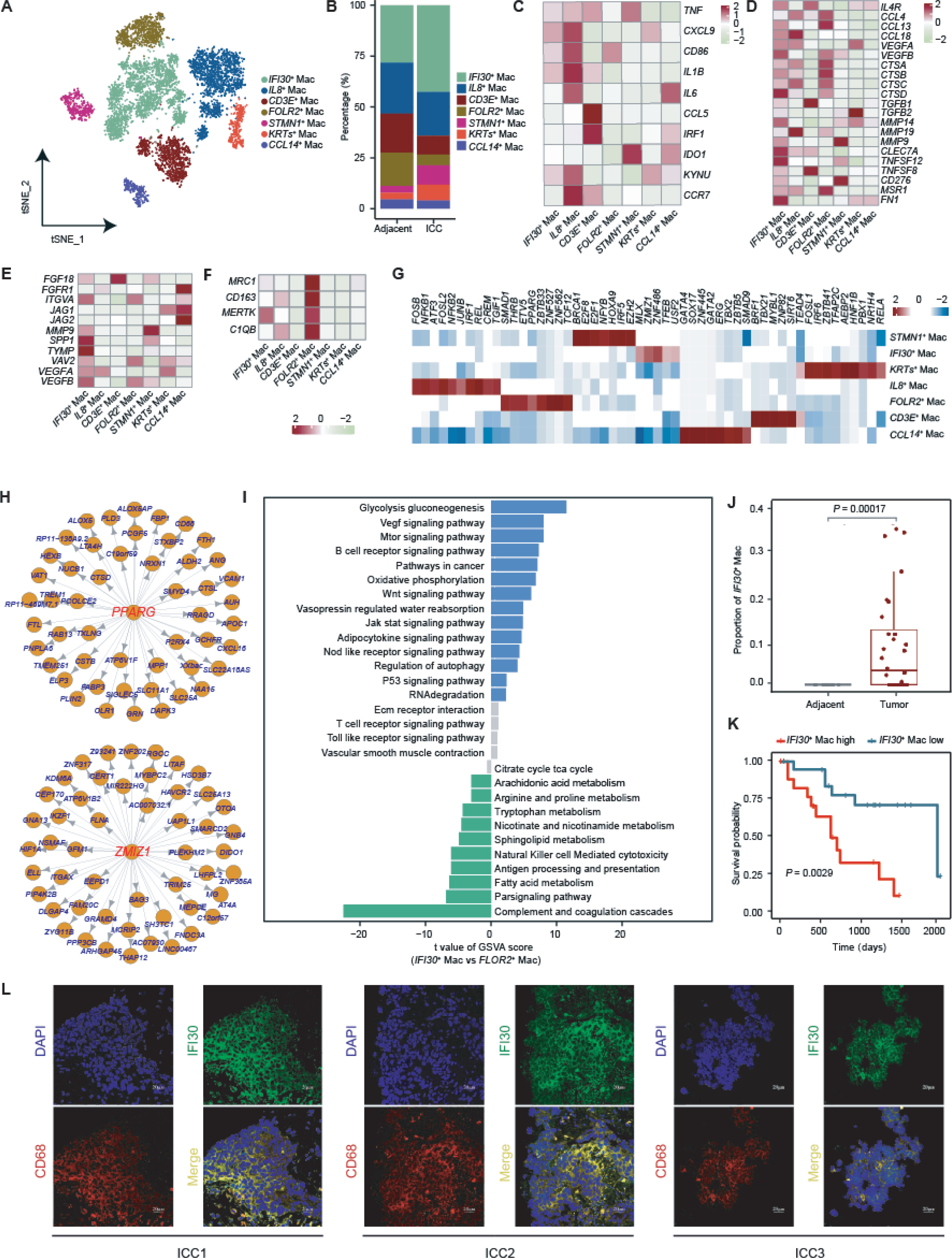
*IFI30*^+^ macrophages are associated with ICC progression. **A.** tSNE plot showing macrophage composition, colored by cluster. **B.** The proportion of macrophage subclusters in adjacent and tumor tissues. **C.** The average expression level of M1 marker genes in macrophage subclusters. **D.** The average expression level of M2 marker genes in macrophage subclusters. **E.** The average expression level of angiogenesis-related marker genes in macrophage subclusters. **F.** The average expression level of phagocytosis-related marker genes in macrophage subclusters. **G.** The enriched transcription factor in macrophage subclusters. **H.** The transcription factors network interacted with PPARG (upper) and ZMIZ1 (lower). **I.** Differences in pathway activity (scored per cell by GSVA) between *IFI30*^+^ and *FLOR2*^+^ macrophages. Blue and green bars represent pathways significant in *IFI30*^+^ macrophages and *FLOR2*^+^ macrophages, respectively. Gray bars represent non-significant pathways. **J.** The differences in infiltration of *IFI30*^+^ macrophages in tumors and adjacent tissues in the TCGA CHOL cohort (n = 50). *P* values were determined by T-test. **K.** Survival analysis of *IFI30*^+^ macrophages in the TCGA CHOL cohort. *P* values were determined by T-test. The mean *IFI30*^+^ macrophages score provided by coxph is used to split samples. Samples with score greater than the mean score of all samples are labeled as “*IFI30*^+^ mac high”, and the others are labeled as *IFI30*^+^ mac low”. **L.** Single channel image for multiplexed immunofluorescence staining of CD68 and IFI30.

To further identify master regulators of *FOLR2*^+^ macrophages and *IFI30*^+^ macrophages, pySCENIC analysis was conducted. The results revealed that the key transcription factor *PPARG* [25, 26], associated with M2 polarization, exhibits higher activity in *FOLR2*^+^ macrophages, further confirming the gradual remodeling of *FOLR2*^+^ macrophages into a pro-tumor phenotype within the ICC tumor microenvironment (Figure 5G, H). GSVA analysis indicated that *FOLR2*^+^ macrophages are predominantly enriched in biological metabolic pathways, including fatty acid metabolism, tryptophan metabolism, and the tricarboxylic acid cycle (Figure 5I). Similarly, the transcription factor *ZMIZ1* [27], known to induce M2 polarization in hepatocellular carcinoma macrophages, displayed high activity in *IFI30*^+^ macrophages, thereby confirming the immunosuppressive phenotype of *IFI30*^+^ macrophages (Figure 5G, H). Genes highly expressed in *IFI30*^+^ macrophages are involved in ECM-receptor interactions, the WNT signaling pathway, and glycolysis/gluconeogenesis. Interestingly, *IFI30*^+^ macrophages are regulated by both the mTOR signaling pathway and the *VEGF* signaling pathway, which are typical hypoxia-induced pathways (Figure 5I). Clinically, CIBERSORTx analysis revealed significantly increased infiltration of *IFI30*^+^ macrophages in tumor tissues (Figure 5J). ICC patients with higher *IFI30*^+^ macrophage abundance exhibited shorter overall survival (OS), a finding validated in the E-MTAB-6389 cohort (Figure 5K, Supplementary Figure 8C). Immunofluorescence confirmed the presence of *IFI30*^+^ macrophages in ICC tissues (Figure 5L).

### *DEFB1*^+^ cholangiocytes and *IFI30*^+^ macrophages co-localized and interacted via TGFB1*/*TGFBR1

Given the involvement of *DEFB1*^+^ cholangiocytes and *IFI30*^+^ macrophages in hypoxia-related pathways, and their association with poor prognosis in ICC patients, we hypothesized potential interactions between these two cell types in hypoxic tumor regions, contributing to an immunosuppressive microenvironment. To explore this, we performed unbiased clustering of spatial transcriptome data from four ICC patients. Although spot clusters varied among patients, *DEFB1*^+^ cholangiocytes and *IFI30*^+^ macrophages consistently co-localized across all samples. They formed a distinct cluster labeled as *DEFB1*^+^ cholangiocytes/*IFI30*^+^ macrophages within tumor regions (Figure 6A, B, and Supplementary Figure 9A-C). The spots of patient ICC1 were divided into 4 clusters, including *DEFB1*^+^ cholangiocytes/*IFI30*^+^ macrophages, cholangiocytes, hepatocytes, and fibroblasts/endothelial cells (Figure 6A, B). The spots of patient ICC2 were divided into 4 clusters, namely *DEFB1*^+^ cholangiocytes/*IFI30*^+^ macrophages, cholangiocytes, hepatocytes/macrophages, and fibroblasts/endothelial cells. The spots of patient ICC3 were divided into 5 clusters, namely *DEFB1*^+^ cholangiocytes/*IFI30*^+^ macrophages, cholangiocytes, hepatocytes, fibroblasts, and endothelial cells/immune cells. The spots of patient ICC4 were divided into 4 clusters, namely *DEFB1*^+^ cholangiocytes/*IFI30*^+^ macrophages, hepatocytes, immune cells, and fibroblasts/endothelial cells (Supplementary Figure 9A-C). We next calculated the scores of the *DEFB1*^+^ cholangiocyte and *IFI30*^+^ macrophage signatures, based on the top 50 specifically expressed genes identified in the scRNA-seq data, in each spot of the spatial transcriptome data. This analysis revealed co-occurrence of these two cell types within the same spatial spots in all patients (Figure 6C, D, Supplementary Figure9 A-C). Additionally, the signature scores of *DEFB1*^+^ cholangiocytes and *IFI30*^+^ macrophages for all spatial spots exhibited a significant positive correlation (Pearson correlation coefficient = 0.88, *P* < 2.2e-16) (Figure 6E, Supplementary Figure 9A-C). These results demonstrated a remarkable co-localization pattern, wherein *DEFB1*^+^ cholangiocytes and *IFI30*^+^ macrophages occupied the same spot, suggesting a physical interaction between these two cell types. Using CIBERSORTx to deconvolute TCGA CHOL tumor samples, we observed a significant positive correlation between *DEFB1*^+^ cholangiocytes and *IFI30*^+^ macrophages (Pearson correlation coefficient = 0.43, *P* = 0.006) (Figure 6F, Supplementary Figure 9D), supporting the interdependence of these two cell types in the context of cholangiocarcinoma. Immunofluorescence further confirmed their spatial proximity (Figure 6G). We investigated the cell-cell interactions in the ICC tumor microenvironment and observed that the number and intensity of ligand-receptor communications between *DEFB1*^+^ cholangiocytes and *IFI30*^+^ macrophages were significantly enhanced compared with para-cancerous tissue (Supplementary Figure 10A, B, Supplementary Table 1-4). Specifically, the activity level of *TGFB1* ligands, which are inferred to regulate *DEFB1*^+^ cholangiocytes by *IFI30*^+^ macrophages, was the most pronounced according to NicheNet (Figure 6H), and the *TGFB1*-encoded protein expressed by *IFI30*^+^ macrophages robustly bound to the *TGFBR1*-encoded receptor present on the surface of *DEFB1*^+^ cholangiocytes (Figure 6I). Notably, spatial transcriptome results showed that *TGFB1* and *TGFBR1* were highly expressed in puncta where *DEFB1*^+^ cholangiocytes and *IFI30*^+^ macrophages co-localized (cell type *DEFB1*^+^ cholangiocytes/*IFI30*^+^ macrophages) (Figure 6J, Supplementary Figure 10C). Further analysis showed that the expression levels of both *TGFB1* and *TGFBR1* in tumor tissues were significantly increased in the TCGA CHOL cohort (Figure 6K). Taken together, these results indicate that the *TGFB1*/*TGFBR1* signaling pathway is significantly enriched in the interaction between *DEFB1*^+^ cholangiocytes and *IFI30*^+^ macrophages.

**Figure 6.**
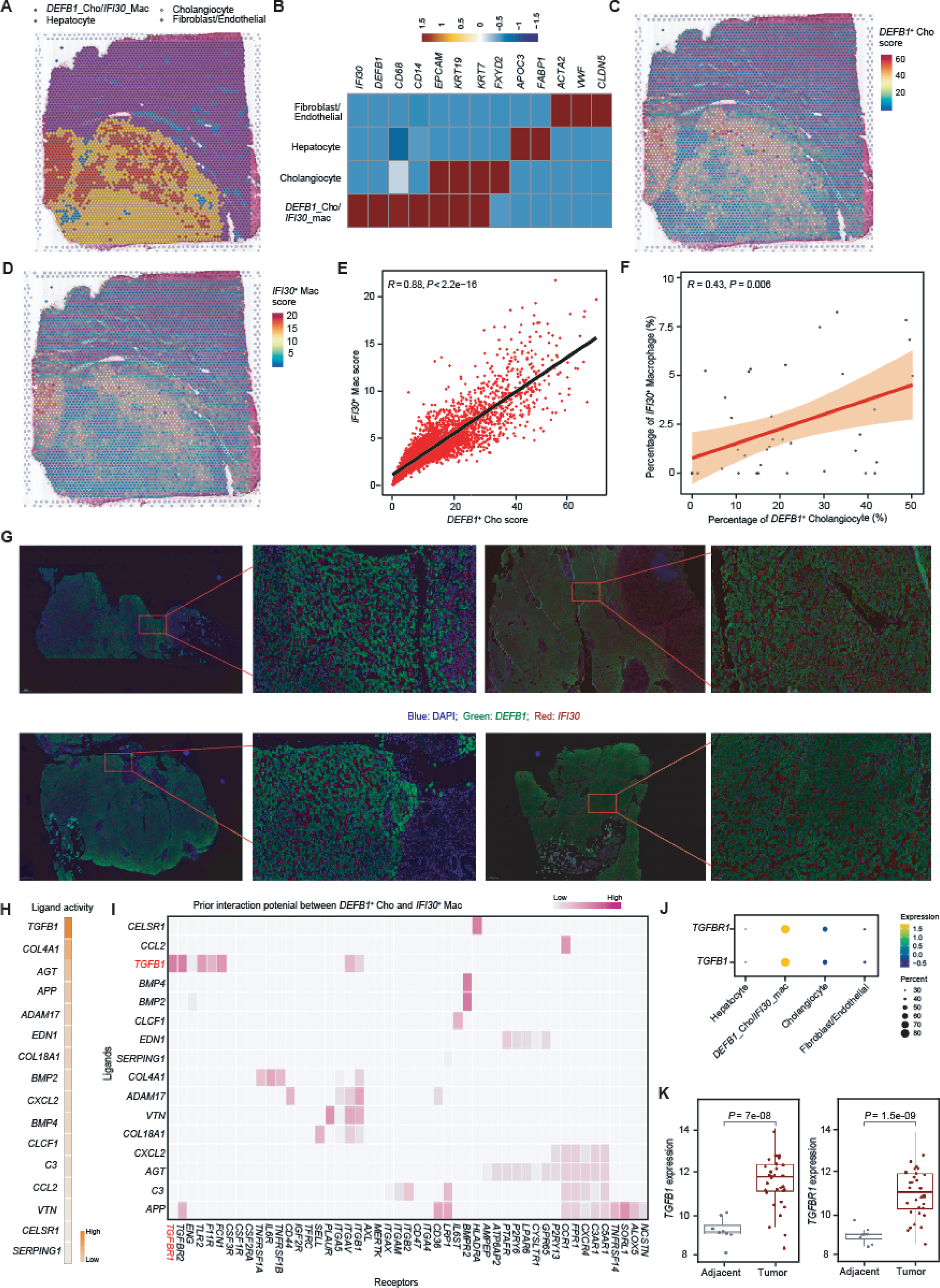
*DEFB1*^+^ cholangiocytes and *IFI30*^+^ macrophages co-localized and interacted via TGFB1/TGFBR1. **A.** Unbiased clustering of spatial spots and definition of cell types in each cluster in the spatial sample ICC1. **B.** Average expression of known markers in indicated cell clusters. **C.** Spatial feature plot of signature score of *DEFB1*^+^ cholangiocytes in the spatial sample ICC1. **D.** Spatial feature plot of signature score of *IFI30*^+^ macrophages in the spatial sample ICC1. **E.** The Pearson correlation of signature score of *DEFB1*^+^ cholangiocytes (x-axis) and IFI30^+^ macrophages (y-axis) in the spatial sample ICC1. *P* values were determined by T-test. **F.** The correlation of infiltration between *DEFB1*^+^ cholangiocytes and *IFI30*^+^ macrophages in the TCGA CHOL cohort. *P* values were determined by T-test. **G.** Single channel image for multiplexed immunofluorescence staining of *DEFB1* and *IFI30*. **H.** Top-ranked ligands inferred to regulate *DEFB1*^+^ cholangiocytes by *IFI30*^+^ macrophages according to NicheNet. **I.** Key ligand-receptor pairs between *DEFB1*^+^ cholangiocytes and *IFI30*+ macrophages. Each row represents a ligand expressed in *DEFB1*^+^ cholangiocytes and each column represents a receptor expressed in *IFI30*^+^ macrophages. **J.** The expression levels of *TGFB1* and *TGFBR1* among cell types in the spatial sample ICC1. **K.** The differences in the expression levels of *TGFB1* (left) and *TGFBR1* (right) between tumor and adjacent tissues in the TCGA CHOL cohort (n = 50). *P* values were determined by T-test.

## Discussion

Immune checkpoint blockade (ICB) has transformed cancer therapy, yet its clinical efficacy in ICC remains limited, with dismal survival outcomes [28, 29]. The high degree of intratumoral heterogeneity coupled with a complex immune microenvironment are key barriers to effective treatment [8]. Our integrated single-cell and spatial transcriptomic analyses provide a comprehensive view of ICC, revealing both cellular diversity and functional interactions that likely shape tumor progression and therapeutic response.

T cells within ICC display marked heterogeneity and dynamic state transitions. Notably, we identified two distinct exhaustion patterns in *CD8*^+^ proliferating T cells, namely terminal exhaustion and progenitor-like exhaustion. These phenotypically and functionally distinct subsets may differentially influence antitumor immunity. Terminally exhausted cells likely reflect irreversible dysfunction, whereas progenitor-like exhausted cells may retain limited proliferative capacity and responsiveness to immunotherapy. Understanding these distinct states is critical for optimizing T cell–directed interventions in ICC.

As the source of malignant tumor cells in ICC, the abnormal status cholangiocytes are crucial for the occurrence and development of ICC [30, 31]. However, currently little is known about the heterogeneity and functional characteristics of cholangiocytes in the tumor microenvironment [32]. Among six identified subpopulations, *DEFB1*^+^ cholangiocytes are positioned at the terminal stage of differentiation, expressing genes linked to angiogenesis, inflammation, and hypoxia adaptation. Their high infiltration correlates with poor overall survival in the TCGA CHOL cohort, suggesting a key role in sustaining tumor growth and shaping the immune microenvironment. These findings highlight *DEFB1*^+^ cholangiocytes as both a prognostic marker and a potential therapeutic target.

Tumor-associated macrophages (TAMs) represent one of the most abundant immune cell types in tumors and can lead to tumor progression, metastasis, and treatment resistance through cross-talk with stromal cells, tumor cells, and other immune cells. Therefore, they have great potential for immunotherapy significance [33]. However, the diversity of TAMs and the mechanisms by which TAM subsets interact with ICC cells remain insufficiently understood. In this study, we systematically characterized the heterogeneity of macrophages and uncovered significant reprogramming of these cells following tumorigenesis. This reprogramming was marked by a specific increase in an immunosuppressive subset (*IFI30*^+^ macrophages) and a reduction in a tissue-resident subset (*FOLR2*^+^ macrophages). Previous research has shown that *FOLR2*^+^ macrophages in breast cancer correlate with better patient survival, likely through enhancing *CD8*^+^ T cell activation [23]. In hepatocellular carcinoma (HCC), however, *FOLR2*^+^ tumor-associated macrophages (TAMs) exhibit an immunosuppressive phenotype, contributing to HCC progression [24]. In our study, *FOLR2*^+^ macrophages in ICC displayed a more pronounced immunosuppressive M2-like gene expression profile, with elevated activity of *PPARG*, a critical transcription factor involved in M2 polarization [34]. This suggests functional similarities between tumor-associated macrophages in ICC and HCC. Meanwhile, *IFI30*^+^ macrophages expressed strong M2-like and pro-angiogenic gene signatures. *ZMIZ1*, a transcription factor known to drive M2 polarization in HCC macrophages [27], was highly active in *IFI30*^+^ macrophages and played a role in hypoxia-induced pathways, including the mTOR and VEGF signaling pathways. Importantly, a higher infiltration of *IFI30*^+^ macrophages was associated with reduced overall survival in ICC patients. Both *DEFB1*^+^ cholangiocytes and *IFI30*^+^ macrophages were found to be closely involved in hypoxia-inducible pathways, and their increased infiltration was linked to poorer patient outcomes. This suggests that hypoxic regions within tumors may exist interactions between *DEFB1*^+^ cholangiocytes and *IFI30*^+^ macrophages, exacerbating the immunosuppressive microenvironment and promoting ICC progression.

Spatial transcriptome results showed that *DEFB1*^+^ cholangiocytes and *IFI30*^+^ macrophages were spatially close, and the infiltration abundance of *IFI30*^+^ macrophages in ICC patients showed a significant positive correlation with the infiltration abundance of *DEFB1*^+^ cholangiocytes, suggesting that the two were more likely to interact. We further used interaction analysis to reveal that the interaction network between *DEFB1*^+^ cholangiocytes and *IFI30*^+^ macrophages in ICC tissues was significantly remodeled, and the *TGFB1*/*TGFBR1* axis was enriched at the interaction site between *DEFB1*^+^ cholangiocytes and *IFI30*^+^ macrophages, while *TGFB1* and *TGFBR1* were both highly expressed at the site where *DEFB1*^+^ cholangiocytes and *IFI30*^+^ macrophages co-localized. Previous research has shown that tumor-associated macrophages can drive cancer stem cell-like properties via *TGFB1*-mediated epithelial-mesenchymal transition (EMT) [35]. In ovarian cancer, macrophage-derived *TGFBI* fosters an immunosuppressive microenvironment, and treatment with anti-*TGFBI* antibodies in mouse models significantly reduced tumor burden [36]. In our study, *TGFB1* and *TGFBR1* expression levels were significantly elevated in tumor tissues compared to adjacent normal tissues, underscoring their critical role in the progression of intrahepatic cholangiocarcinoma (ICC).

Collectively, our findings have several translational implications. First, strategies aimed at modulating *DEFB1*^+^ cholangiocytes or *IFI30*^+^ macrophages may disrupt tumor-supportive interactions. Second, targeting the *TGFB1*/*TGFBR1* axis offers a potential combinatorial approach with existing immunotherapies. Third, the heterogeneity within *CD8*^+^ T cells suggests that interventions may need to account for both terminally and progenitor-exhausted populations to maximize efficacy.

Limitations of this study include the partial pairing of spatial and single-cell datasets from the same patient, which may obscure inter-patient heterogeneity, and the need for mechanistic studies to validate *TGFB1*/*TGFBR1*–mediated interactions. Despite these constraints, our work provides a high-resolution atlas of the ICC microenvironment, integrating immune and non-immune cell populations, spatial organization, and functional interactions. These insights lay a foundation for precision-targeted therapies aimed at overcoming immune suppression and improving outcomes for ICC patients.

## Materials and Methods

### Sample and data collection

With informed consent and approval from the local Medical Ethics Committee, normal, paracancerous, and tumor tissues were collected from an intrahepatic cholangiocarcinoma (ICC) patient who was confirmed by clinical pathological diagnosis at the Affiliated Hospital of Guizhou Medical University for sequencing. Additionally, 49 paraffin sections from 34 patients were obtained for immunohistochemistry and immunofluorescence experiments.

Single-cell transcriptome data of 16 intrahepatic cholangiocarcinoma samples, including 11 tumor tissues and 5 adjacent tissues, were downloaded from GEO (GSE138709 and GSE142784) and GSA-human (HRA001748). Spatial transcriptome data of 3 intrahepatic cholangiocarcinoma samples were also collected from GSA-human (HRA002304 and HRA000437). Both single-cell and spatial transcriptome data were generated using the 10x Genomics platform. For survival analysis, we further incorporated two publicly available transcriptomic cohorts, including TCGA-CHOL (n = 50) and E-MTAB-6389 (n = 105).

### Single-cell transcriptome sequencing and bioinformatics analysis Single-cell transcriptome sequencing

The SIC single-cell suspension (700-1200 cells per ml as determined by CellDrop FL cell counter) was prepared using the Chromium Single Cell 3’ GEM, Library & Gel Bead Kit v3.1 (10x Genomics, PN-1000268) according to the manufacturer’s instructions for live cells. Subsequently, the suspension was loaded onto a Chromium Single Cell Chip (Chromium Single Cell G Chip Kit, 10x Genomics, PN-1000120) and co-encapsulated with barcoded gel beads to achieve a target capture rate of 6000 cells per sample. The captured cells were then lysed, and the released RNA (GEMS) was barcoded through reverse transcription of individual single-cell gel beads within emulsion. Within each droplet, complementary DNA (cDNA) was generated and amplified by reverse transcription on a T100 PCR thermal cycler (Bio-Rad), with reaction conditions set at 53°C for 45 minutes, followed by 85°C for 5 minutes, and holding at 4°C. Subsequently, the concentration and quality of the cDNA were assessed using a Qubit fluorometer (Thermo Scientific) and a bioanalyzer 2100 (Agilent), respectively. According to the manufacturer, the scRNA-seq library was constructed and sequenced on the Illumina platform (novaseq 6000) by Annouda Gene Technology Co., Ltd, Beijing, China, with a depth of 40,000 reads per cell.

### Data preprocessing

The CellRanger software pipeline (version 5.0.0, 10x Genomics) was employed for raw sequencing read alignment, annotation, and quantification (utilizing the genome reference set GRCh38-3.0.0). The unique molecular identifier (UMI) count matrix was processed using the R package Seurat (version 4.0.5) [37], with cells expressing fewer than 200 genes and comprising more than 20% mitochondrial reads being excluded. Potential doublets were identified and filtered out using the software package DoubletFinder (version 2.0.3) [38]. The scRNA-seq data were integrated using canonical correlation analysis (CCA) algorithms to address batch effects among samples.

### Dimension reduction and clustering analysis

Library size normalization was conducted using the NormalizeData function in Seurat to obtain normalized counts, followed by log transformation of the results. The top 2000 highly variable genes (HVGs) were identified utilizing the FindVariableFratures function. Dimensionality reduction of the data based on HVGs was performed through principal component analysis (PCA). To generate a two-dimensional representation of cell states, the matrix underwent uniform manifold approximation and projection (UMAP) dimensionality reduction analysis. The clustering of cells was carried out using the FindClusters function, employing a method based on shared nearest neighbor module optimization.

### Differentially expressed gene identification and cell type annotation

The FindAllMarker function in Seurat was utilized to perform a Wilcoxon rank-sum test, aiming to identify differentially expressed genes in each cluster. For the particular cluster, the FindAllMarkers function identifies positive markers in comparison to all other clusters. Default values were applied to all parameters. Cell types were manually annotated by integrating differentially expressed genes with cell type marker genes reported in the literature.

### Copy number variation analysis

The infercnv package [39] was employed to identify copy number variations (CNVs) in cholangiocytes. The subset function was utilized to extract the gene expression matrix of reference cells (including dendritic cells, macrophages, and B cells) and cholangiocytes. Subsequently, the CreateInfercnvObject function was utilized to create a CNV analysis object, and the infercnv::run function was applied to identify the copy number variation levels of cholangiocytes.

### Pseudo time analysis

Developmental trajectories of cholangiocytes and T cells were inferred utilizing Monocle3 (version 1.3.4) [40]. The gene expression matrix of a specific cell type was extracted into Monocle3 to construct a CellDataSet object using the subset function. Subsequently, after preprocessing and dimensionality reduction of the analysis objects through the preprocess_cds and reduce_dimension functions, respectively, a series of representative key-acting genes were unveiled along the differentiation process via the graph_test function. The visualization function plot_cells was employed to depict each group along the same pseudo-time trajectory.

### Transcription factor regulon analysis

Gene regulatory networks and regulon activities were inferred using pySCENIC [41]. Regulon activity, measured as area under the curve (AUC), was analyzed utilizing pySCENIC’s AUCell module, with active regulons determined based on default thresholds set by AUCell. Differential-expression regulons were identified via the Wilcoxon rank-sum test in the FindAllMarkers function, with all parameters set to default values. Regulon activity heatmaps were generated using the Pheatmap package [42].

### GSVA analysis and functional enrichment analysis

Enrichment analysis of Hallmark gene sets was conducted utilizing the GSVA package [43] and msigdbr package [44] to investigate tumor-related biological states and processes in the most significantly enriched cell types. Additionally, the top 100 differentially expressed genes of each cell type were extracted, and functional enrichment analysis was performed using the DAVID online website (https://david.ncifcrf.gov/tools.jsp).

### Characterization of cell-type infiltration based on single-cell expression matrix

To determine the proportions of our annotated 14 major cell types and 38 cell subtypes from RNA-seq data, we utilized the online tool CIBERSORTx [45] to generate a reference signature matrix from our single-cell RNA-seq dataset and to estimate cell-type proportions from the TCGA cholangiocarcinoma (CHOL) cohort based on the constructed cell-type reference (https://portal.gdc.cancer.gov). All parameters were set to default values. Pearson correlation analysis was employed to assess the relationship between cell type infiltration proportions, with correlations exceeding |R| > 0.3 and FDR < 0.05 considered significant. Unsupervised clustering using the R package Pheatmap was employed to evaluate and visualize the level of correlation of cellular infiltration among all cell types in the TCGA CHOL cohort.

### Survival analysis

All tumor samples were divided into two groups, namely high infiltration and low infiltration, based on the median infiltration abundance. Survival curves were plotted using the Kaplan-Meier method and the Survival software package (version 2.44), and visualization was carried out utilizing the ggsurvplot function of the survminer software package. Significance was assessed via the log-rank test statistic (p-value) between the two groups [46].

### Cell–cell communications analysis by CellChat

The normalized gene expression matrix and cell type annotation results were extracted using the Subset function, and an analysis object was created using the createCellChat function in the CellChat package [47]. Overexpressed ligands or receptors in cells were identified using the determineOverExpressedGenes function, and the determineOverExpressedInteractions function utilized overexpressed ligands or receptors to construct protein-protein interaction (PPI) networks. The computeCommunProb function was then used to model inter-cell communication probability and predict cell communication networks.

### Ligand receptor analysis by NicheNet

The NicheNet package [48] was employed to infer ligand-receptor activity between *DEFB1*^+^ cholangiocytes and *IFI30*^+^ macrophages. For ligand and receptor interactions, genes expressed in more than 10% of cluster cells were considered. The top 100 ligands and the top 1000 targets of differentially expressed genes in “sending cells” (*IFI30*^+^ macrophages) and “affected cells” (*DEFB1*^+^ cholangiocytes) were extracted for paired ligand-receptor activity analysis. The Nichenet_output$ligand_activity_target_heatmap function was utilized to plot ligand-modulated activity. Activity scores range from 0 to 1.

### Spatial transcriptome sequencing and bioinformatics analysis Spatial transcriptome sequencing

A fresh ICC tissue sample was surgically excised and embedded in pre-cooled OCT in a petri dish. The embedded tissue and cassette were then transferred to dry ice powder and frozen for half an hour until the OCT was completely frozen. Subsequently, the cassette was placed on dry ice to proceed with the operation. The operational steps were as follows: (1) Tissue optimization: The tissue sections were fixed using methanol and stained with hematoxylin and eosin. Following staining, the tissue was fixed, stained, and permeabilized to release mRNA. (2) Expression library construction: mRNA was captured, and a reverse transcription reaction was performed to obtain full-length cDNA labeled with Spatial Barcode. A second-strand reaction mix was added to synthesize the second strand, followed by cDNA denaturation, transfer, recovery, and amplification. (3) Library construction: The captured RNA was used as a template to prepare a cDNA synthesis and sequencing library. (4) Library quality inspection: The fragment length distribution of the library was detected using Agilent 2100/Lab Chip GX Touch. Once the library met expectations, Q-PCR was employed to accurately quantify the effective concentration of the library (with an effective concentration >10 nmol/L) to ensure library quality. (5) Sequencing: Qualified libraries were tested and sequenced on the Illumina HiSeq/Novaseq sequencing platform.

### Spatial transcriptome data analysis

Space Ranger (version 1.3, 10x Genomics) was utilized to align the raw sequencing reads with the human GRCh38 reference genome and parse the spatial position information. The spatial transcriptome data were processed using Seurat (version 4.0.5) [37] to generate a gene expression matrix for analysis. To mitigate batch effects, methods similar to those used for single-cell transcriptome data analysis were employed, including the FindIntegrationAnchors, SCTransform, and IntegrateData functions. Dimension reduction of integrated data was performed using PCA. The uniform manifold approximation and projection (UMAP) method was applied to visualize a 2D projection of cell clusters from an SNN graph. The FindClusters function was used to cluster spots. To predict the main cell type at each spot, the RCTD package [49] was used to assign a mixture of single-cell transcriptome annotated cell types to spatial transcriptome spots, with the mode selected as “full mode”. The plotSpatialScatterpie function in the SPOTlight package [50] was utilized to visualize cell type deconvolution results, and the plotInteractions function was employed to conduct spatial co-localization analysis.

### Immunohistochemistry

Paraffin sections were heated in a 70°C oven for 30 minutes, followed by deparaffinization and antigen retrieval. To assess the expression patterns of candidate antibodies in ICC and adjacent tissues, primary antibodies were applied and incubated overnight at 4°C. Secondary antibodies were used for immunostaining, with hematoxylin used for counterstaining. Antibody details: *CD3* (Catalog No.: 60181, Proteintech), CD68 (Catalog No.: AB955, Abcam).

### Immunofluorescence Staining

Place paraffin sections in a 70°C oven for 30 minutes. Deparaffinize the slides, then immerse in PBS briefly. Add diluted antigen retrieval solution to the staining incubation box, place the slides inside, cover, and apply high pressure at 120°C for 2 minutes. After cooling to room temperature, discard the solution, immerse the slides in PBS for 2 minutes, wash three times, and wipe off the PBS around the tissue. Add 300 μl of sheep serum to cover the tissue, incubate at 37°C for 30 minutes, then discard the blocking solution and wash with PBS for 2 minutes. Dilute the primary antibody at 1:100, apply 100 μl per slide, and incubate overnight at 4°C. The next day, wash the slides three times in PBS for 2 minutes each. Add diluted secondary antibody (1:200) in the dark and incubate at room temperature for 30 minutes. Wash three times in PBS, apply 100 μl of DAPI solution per slide, incubate at room temperature for 5 minutes, and wash again. Dry the tissue, then mount using anti-fading fluorescence medium. Antibody details: CD68 (1:500, Abcam, AB213363), EPCAM (1:100, Abcam, AB7504), DEFB1 (1:100, Abbexa, abx109572), IFI30 (1:100, LSBio, LS-C757291).

## Data availability

The raw sequence data reported in this paper have been deposited in the Genome Sequence Archive of the BIG Data Center at the Beijing Institute of Genomics, Chinese Academy of Science, under accession number HRA010008 (accessible at http://bigd.big.ac.cn/gsa-human). The processed ICC public single cell transcriptome dataset were downloaded from Gene Expression Omnibus (https://www.ncbi.nlm.nih.gov/geo/, GSE138709 and GSE142784) and GSA-human (https://ngdc.cncb.ac.cn/gsa-human/, HRA001748). The processed ICC public Spatial transcriptome dataset were downloaded from GSA-human (https://ngdc.cncb.ac.cn/gsa-human/, HRA002304 and HRA000437). Classic transcriptome datasets for survival were downloaded from TCGA (https://www.cancer.gov/ccg/research/genome-sequencing/tcga, TCGA CHOL cohort) and EMBL-EBI (https://www.ebi.ac.uk/, E-MTAB-6389).

## CRediT authorship statement

**Guoliang Wang:** Investigation, Data curation, Formal analysis, Validation, Visualization, Writing – original draft, Writing – review & editing. **Hang Meng:** Formal analysis, Validation, Writing – original draft, Writing – review & editing. **Meng Li:** Formal analysis. **Xiangdong Fang:** Conceptualization, Resources, Writing – original draft, Writing – review & editing, Supervision. **Weilong Zou:** Conceptualization, Resources, Writing – original draft, Writing – review & editing, Supervision. **Hongzhu Qu:** Conceptualization, Resources, Writing – original draft, Writing – review & editing, Supervision.

## Competing interests

The authors have declared no competing interests.

## Acknowledgments

This research was supported by the Strategic Priority Research Program of the Chinese Academy of Sciences (grant no. XDA0460403) and Doctoral Research Foundation Project of the Affiliated Hospital of Guizhou Medical University (grant no. gyfybsky-2022-1).

## Supplementary material

**Figure S1.**
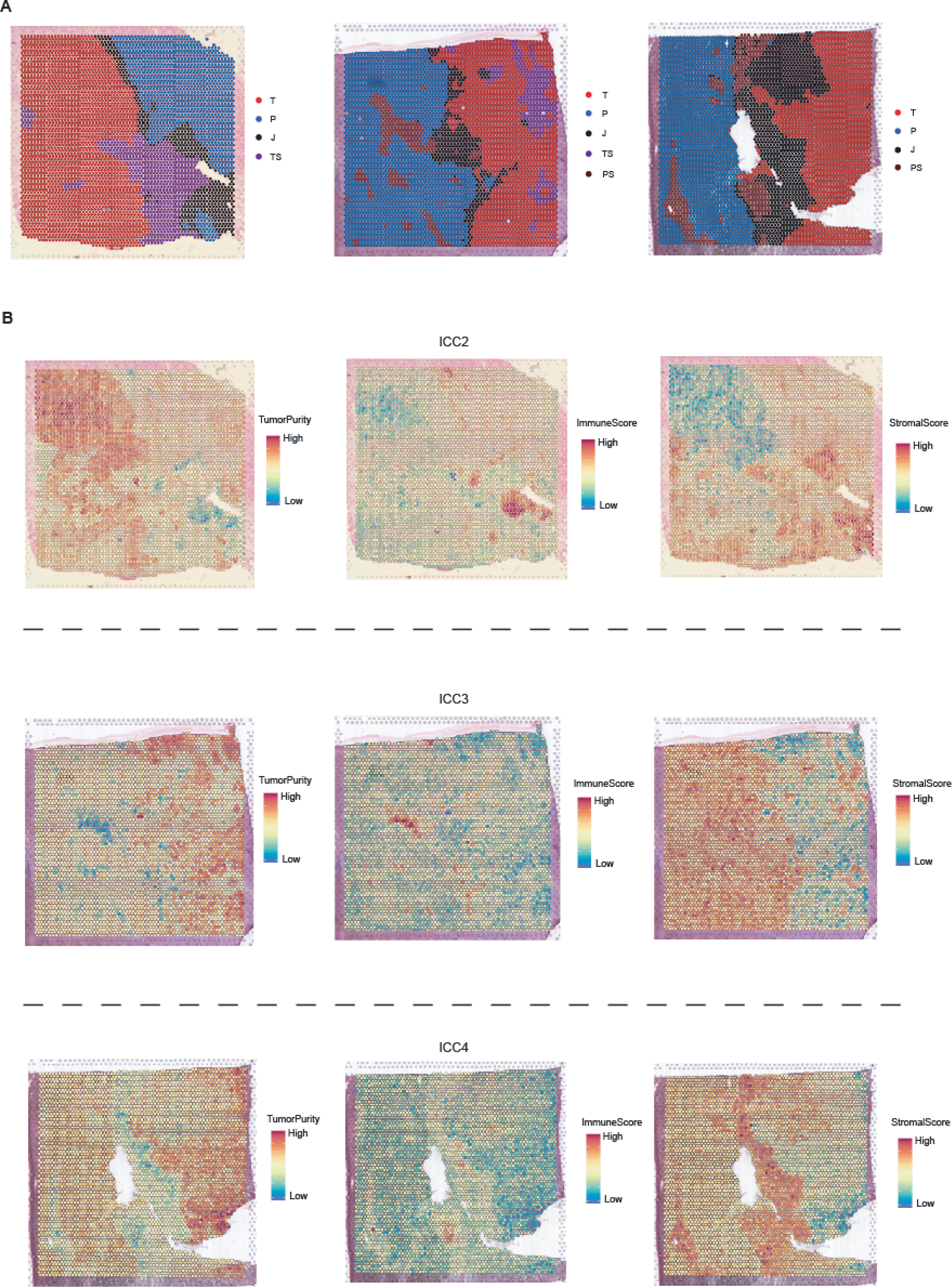
Spatial transcriptome heterogeneity of ICC. **A.** Pathologists manually annotated the tumor area, adjacent area, invasion area, tumor stroma, and adjacent stroma in the paraffin section of patients (patient 2, patient 3 and patients 3). J, invasive front region. P, adjacent region. T, tumor region. TS, tumor stroma region. PS, adjacent stroma region. **B.** Expression data for each spatial feature was assessed using ESTIMATE to generate stromal score, immune score and tumor purity score in the spatial sample ICC2 (top), ICC3 (medium), and ICC4 (bottom), respectively.

**Figure S2.**
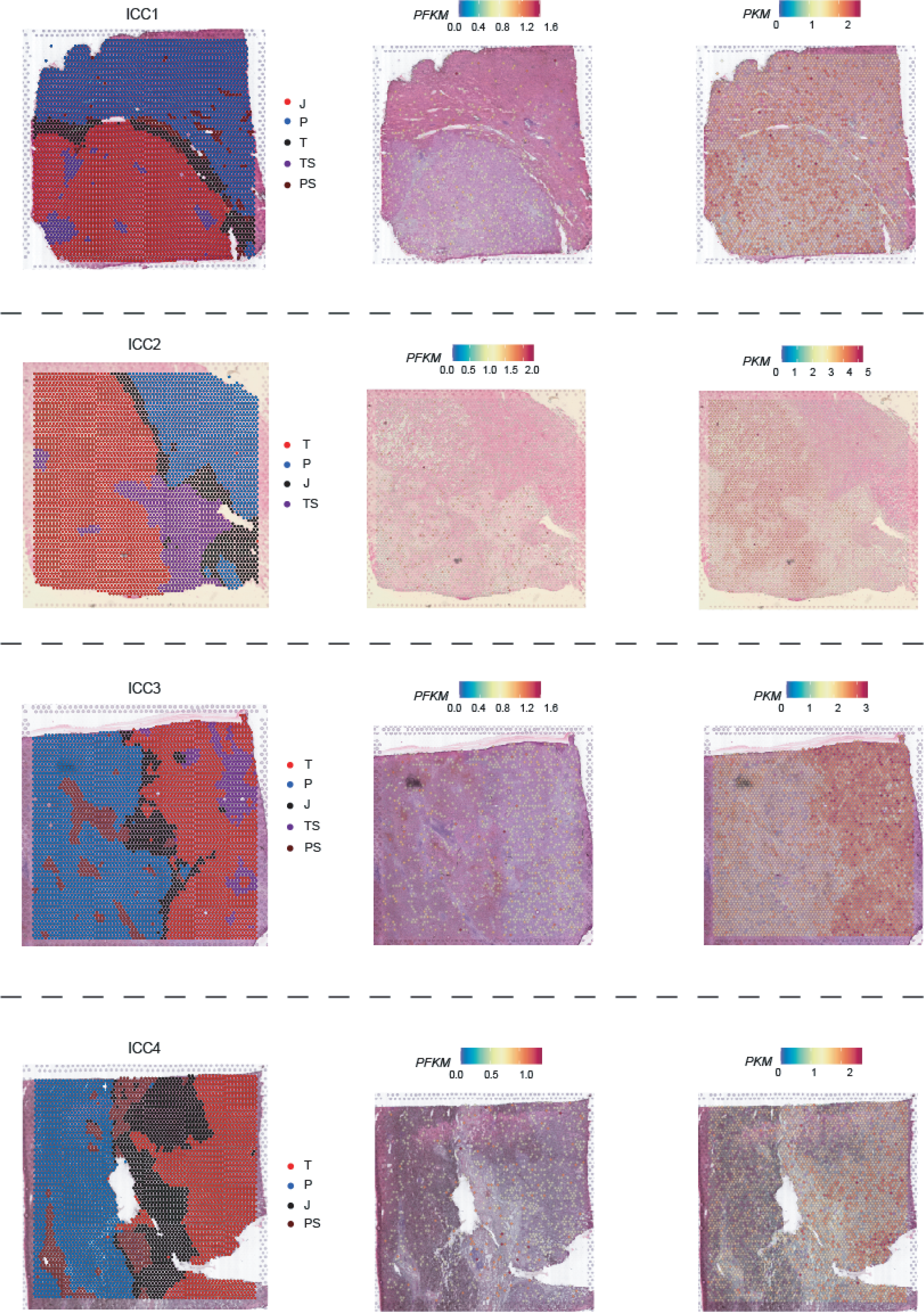
Spatial expression patterns of key genes in glycolysis. Spatial expression patterns of key glycolysis genes (*PFKM* and *PKM*) in four ICC patients

**Figure S3.**
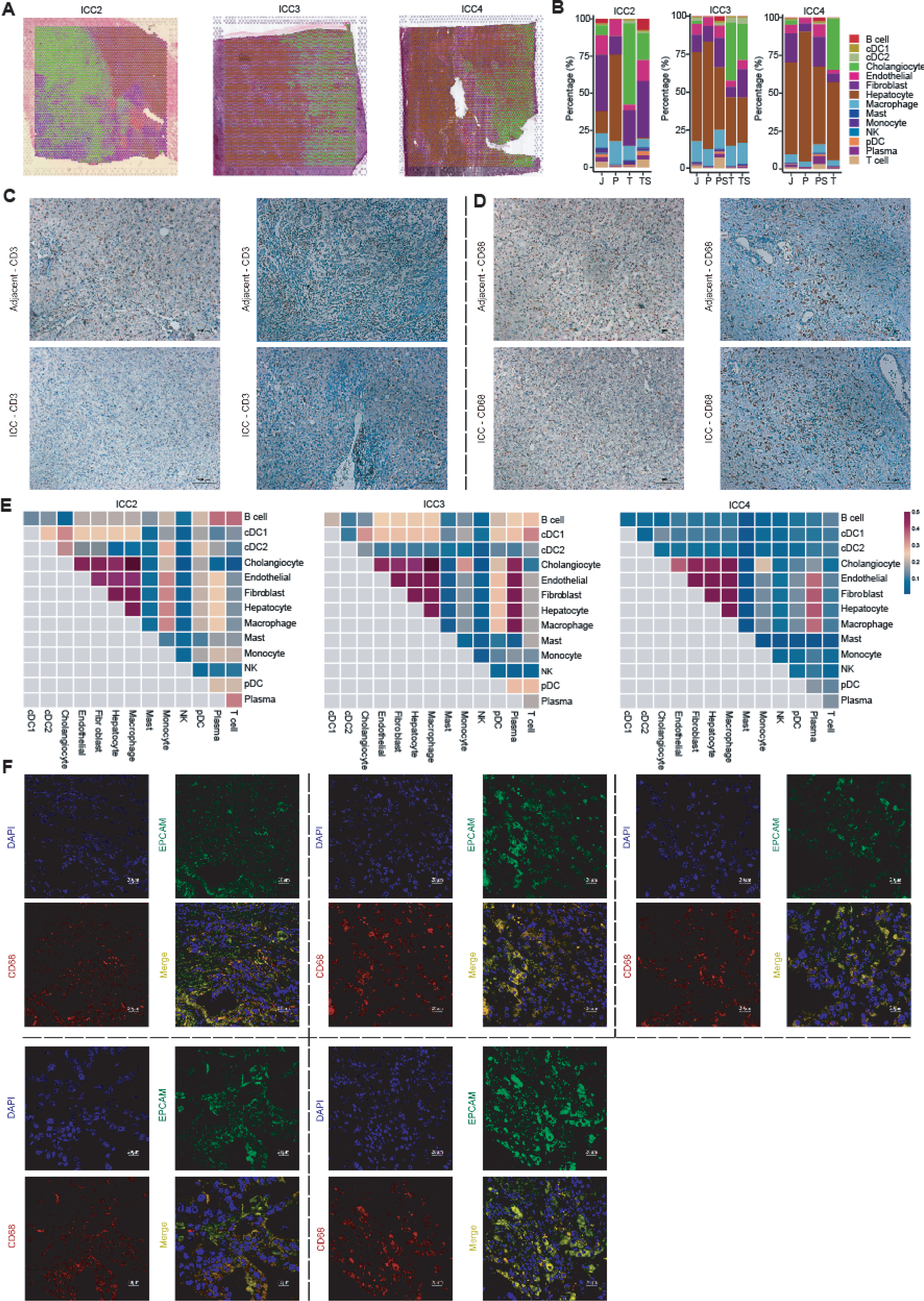
Spatial transcriptome heterogeneity of ICC. **A.** Deconvolution analysis of cell types in spatial samples ICC2, ICC3, and ICC4, based on single-cell transcriptome annotations. **B.** The bar chart shows the proportions of different cell types in various tissue regions of the spatial samples. **C.** The expression of *CD3* (T cells marker) in ICC tissue and adjacent tissue by the immunohistochemical examination. **D.** The expression of *CD68* (macrophages marker) in ICC tissue and adjacent tissue by the immunohistochemical examination. **E.** The spatial co-localization levels of paired cell types in spatial samples (ICC2, ICC3, and ICC4). **F.** Single channel image for multiplexed immunofluorescence staining of *EPCAM* and *CD68*.

**Figure S4.**
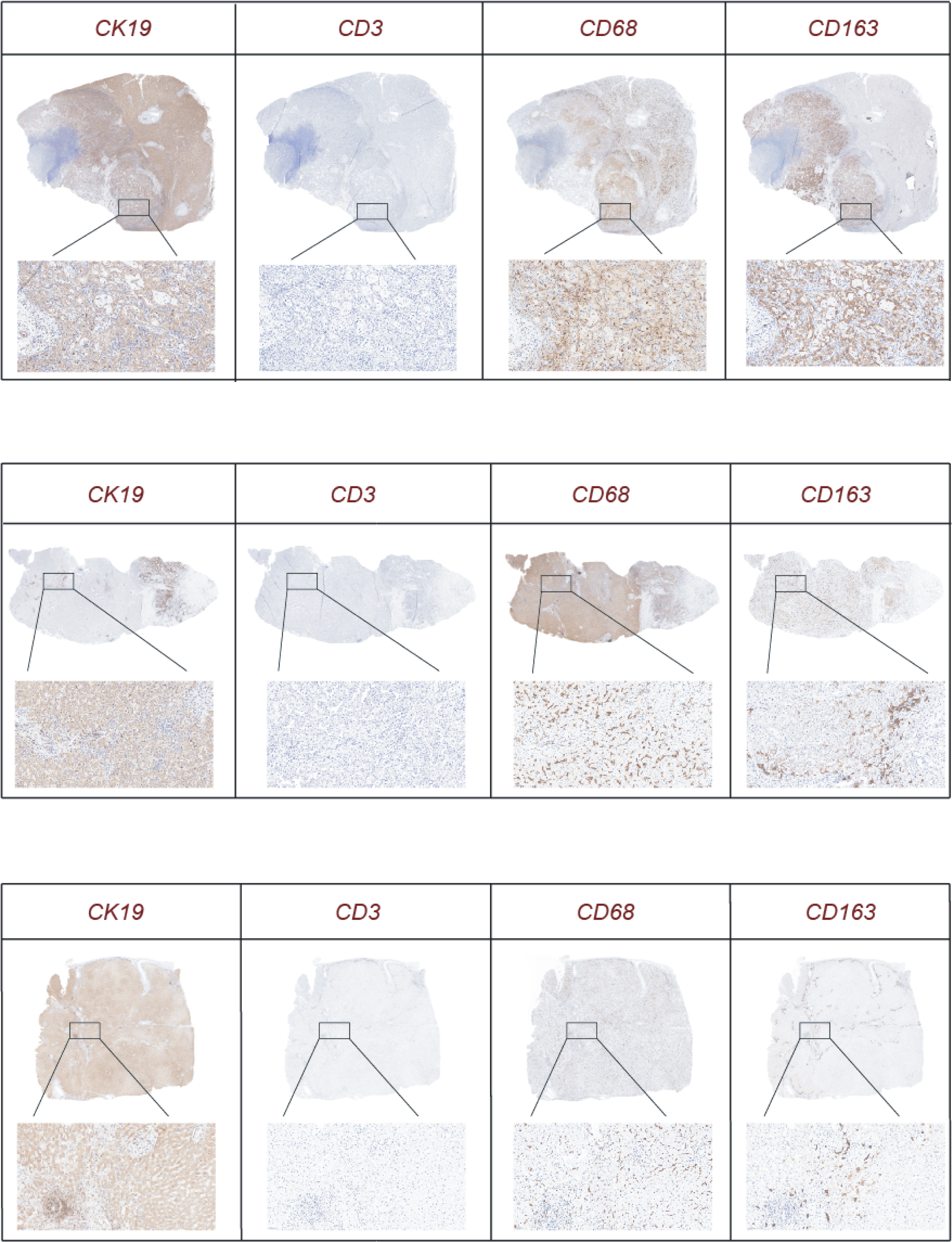
Immunohistochemical detection of key proteins in ICC tissue. Immunohistochemical detection of *CK19* (tumor cell marker), *CD3* (T cell marker), *CD68* (macrophage marker), and *CD163* (M2 macrophage marker) in ICC tissue.

**Figure S5.**
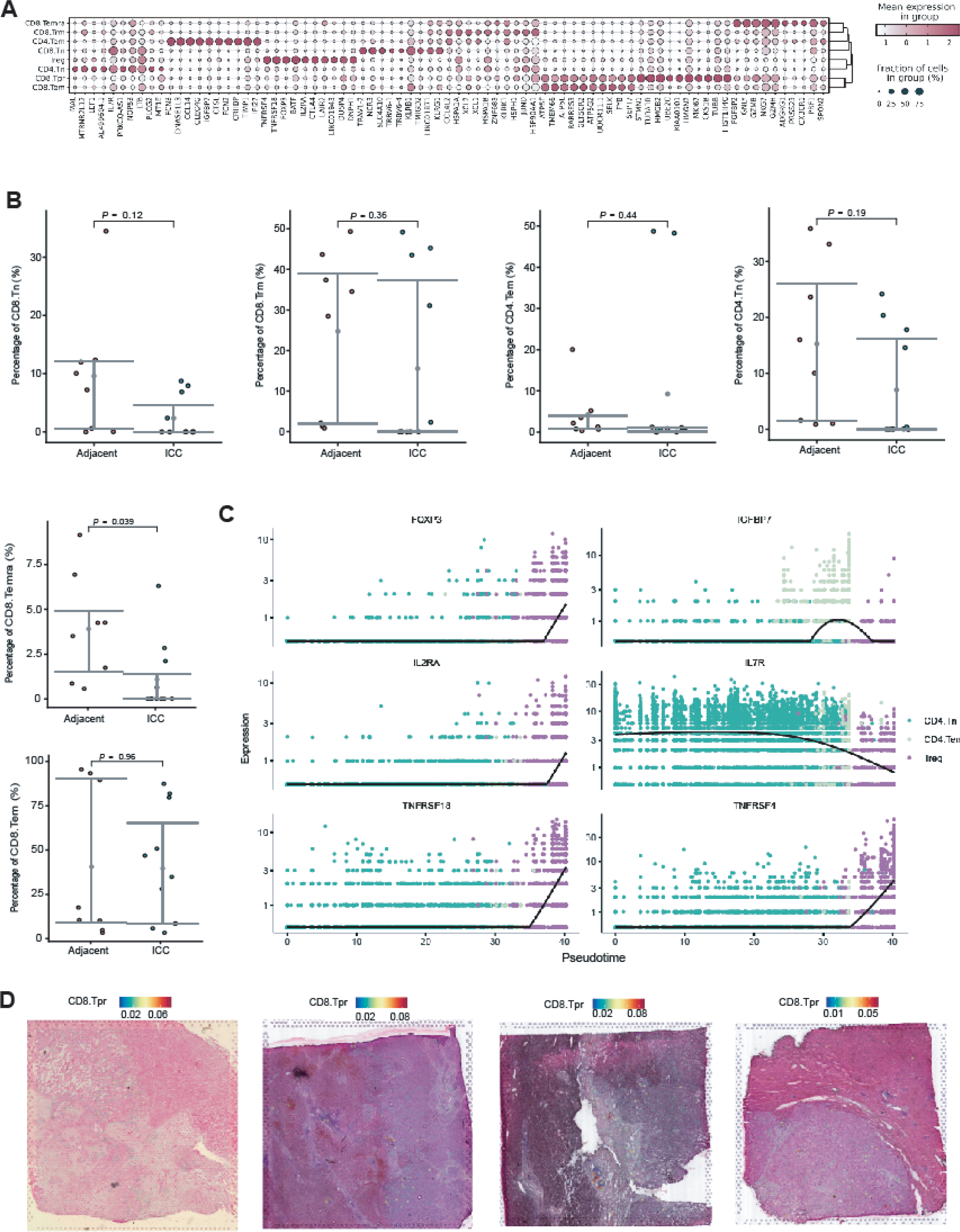
Heterogeneous exhaustion patterns of *CD8*^+^ Tpr populations in ICC patients. **A.** The top-ranked differentially expressed genes (DEGs) in T cell subclusters. **B.** The differences in proportions of T cell subclusters between peritumoral and tumor tissues. *P* values were determined by T-test. **C.** Visualization of the top six DEGs along the pseudotime trajectory selected according to Moran’s I index in the three *CD4*^+^ T cell subclusters. **D.** Distribution of *CD8*^+^ proliferating T cells in the four spatial samples.

**Figure S6.**
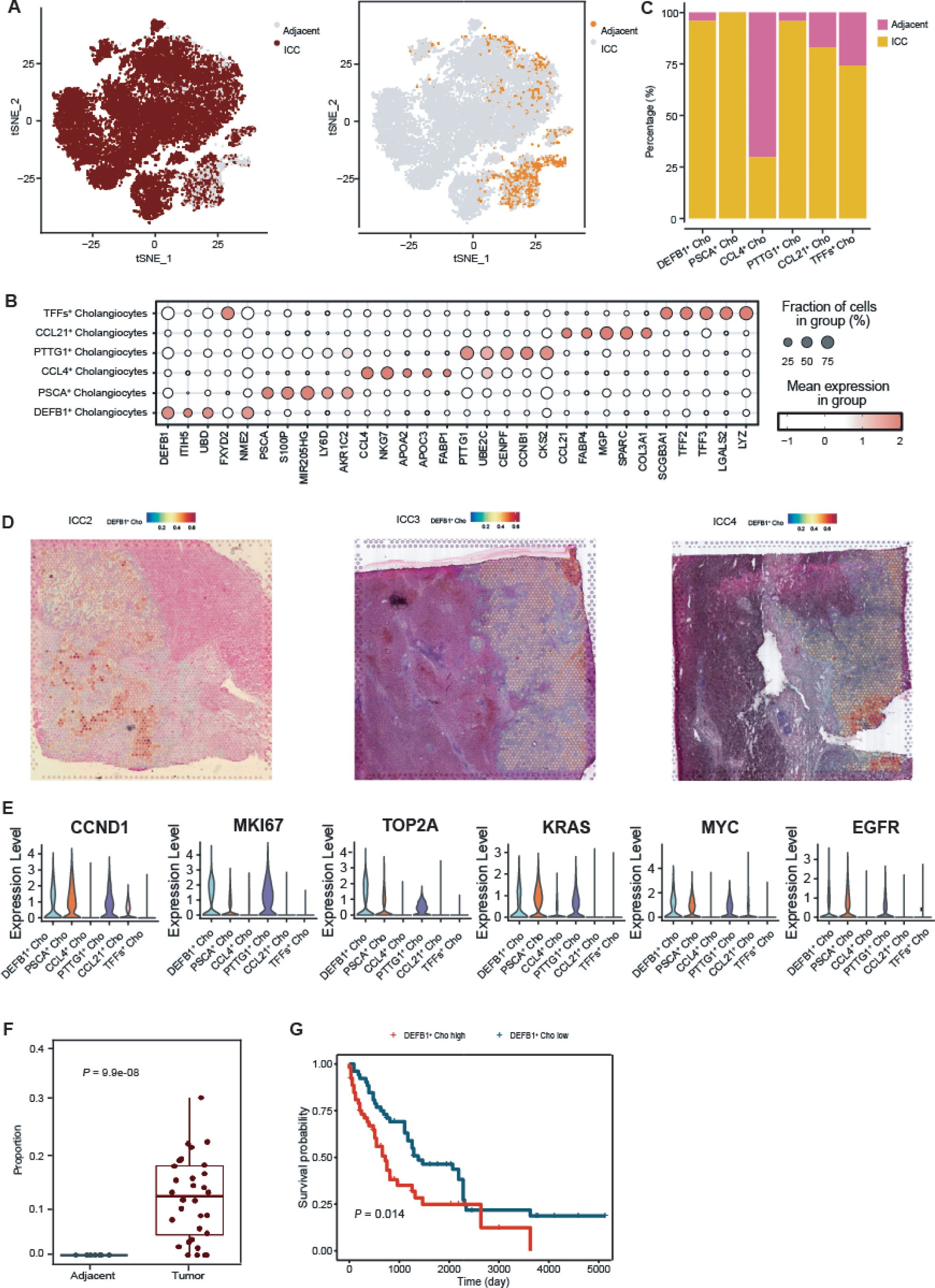
*DEBF1^+^* malignant cells are associated with ICC progression. **A.** tSNE plot demonstrating the distribution of cholangiocytes in tumor and adjacent tissues. **B.** The top-ranked DEGs in cholangiocyte subclusters. **C.** The distribution of cholangiocyte subtypes in tumor tissue and adjacent tissues. **D.** Distribution of *DEFB1*^+^ cholangiocytes in spatial samples (ICC2, ICC3 and ICC4). **E.** Expression levels of oncogenes (*KRAS*, *MYC*, *CCND1*, and *EGFR*) and proliferation-related genes (*MKI67* and *TOP2A*) in cholangiocyte subtypes. **F.** The difference in infiltration of *DEFB1*^+^ cholangiocytes between tumor and adjacent tissues in the TCGA CHOL cohort (n = 50). *P* values were determined by T-test. **G.** Survival analysis of *DEFB1*+ cholangiocytes in the E-MTAB-6389 cohort (n = 105). *P* values were determined by T-test. The mean *DEFB1*^+^ cholangiocytes score provided by coxph is used to split samples. Samples with score greater than the mean score of all samples are labeled as “*DEFB1*^+^ cho high”, and the others are labeled as *DEFB1*^+^ cho low”.

**Figure S7.**
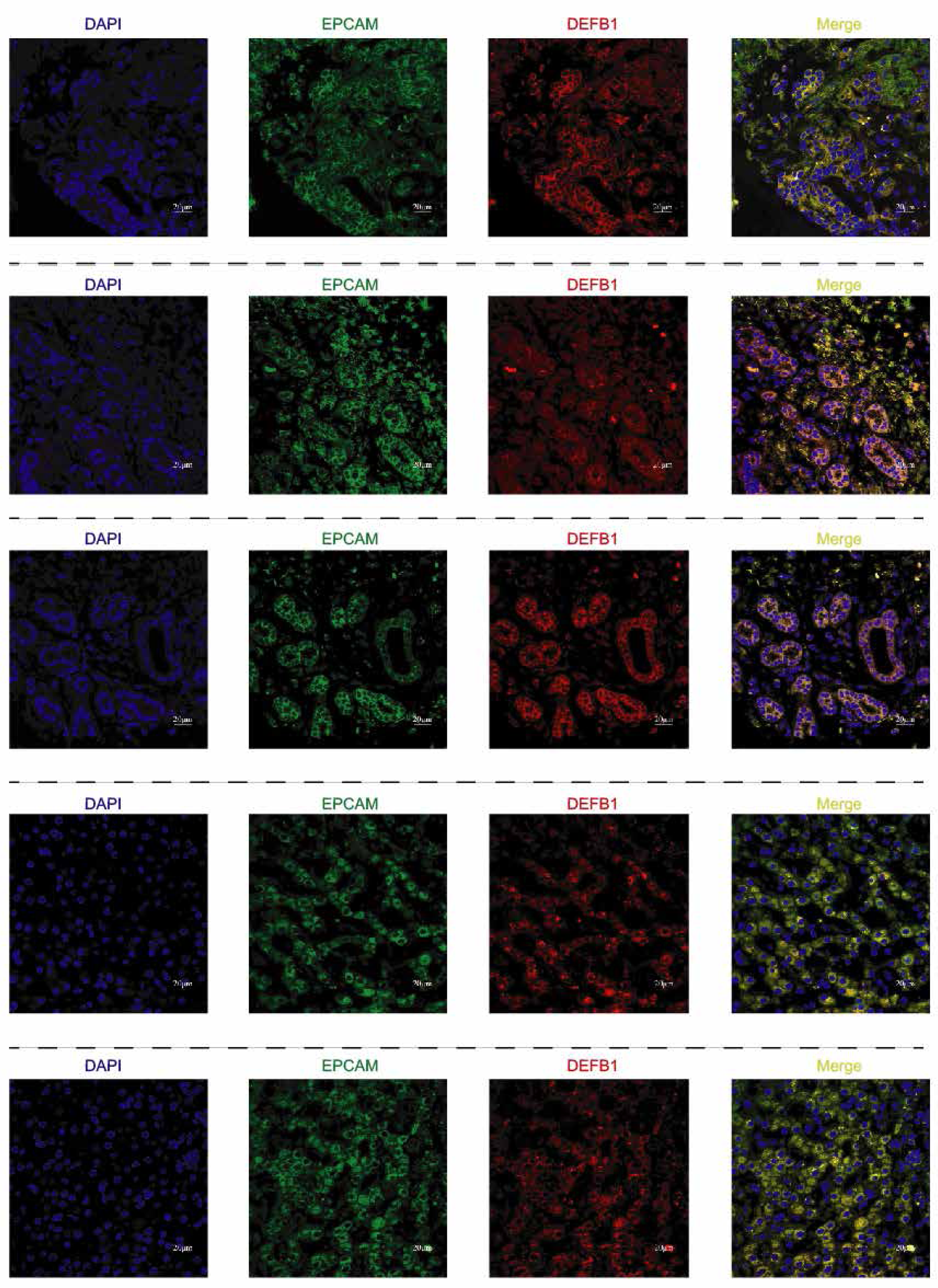
*DEBF1*^+^ malignant cells are associated with ICC progression. Single channel image for multiplexed immunofluorescence staining of *EPCAM* and *DEFB1* detected in the tumor sections of 5 validation patients.

**Figure S8.**
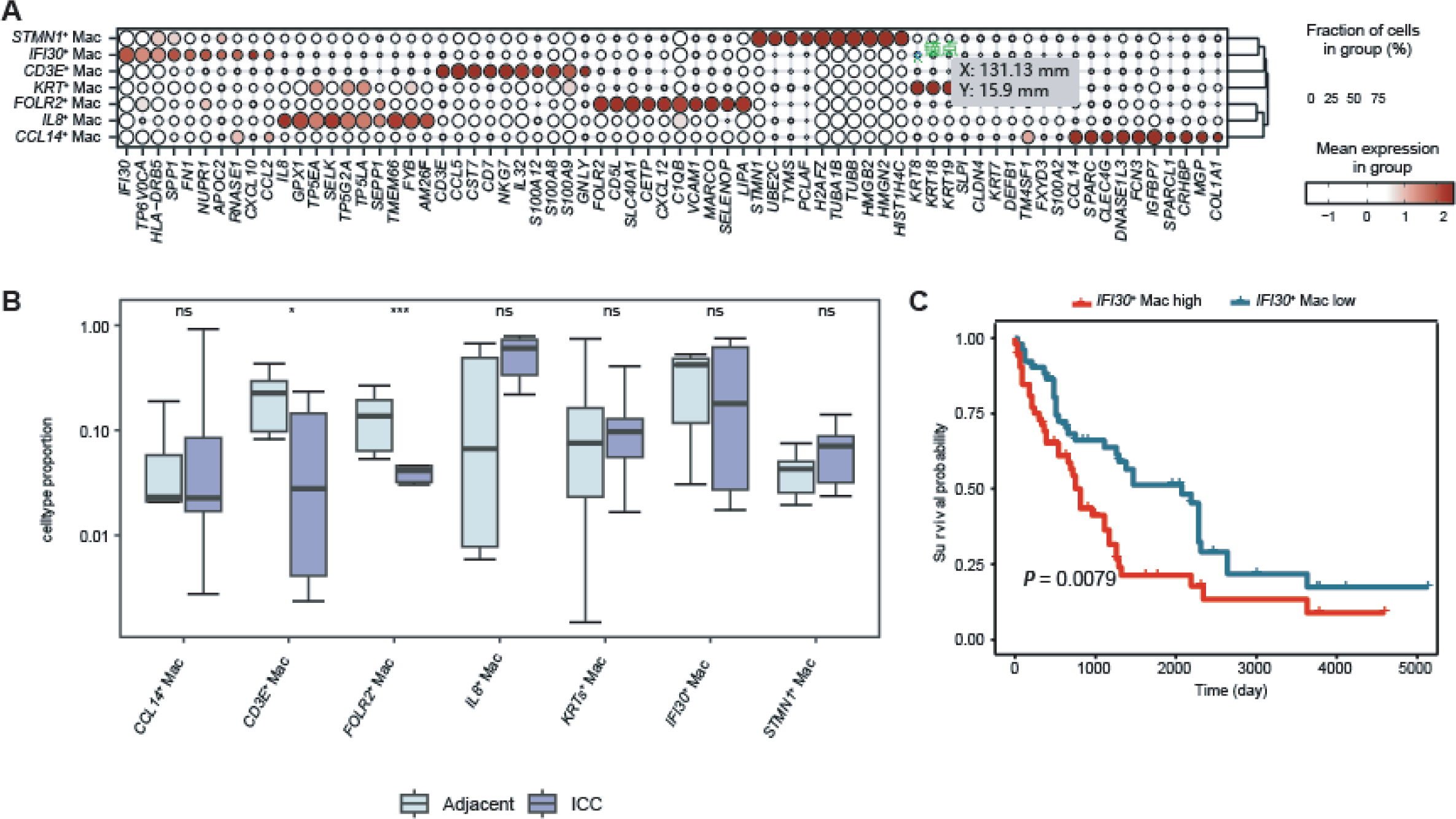
*IFI30*^+^ macrophages are associated with ICC progression. **A.** The top-ranked DEGs in macrophage subclusters. **B.** The differences in the proportions of different cell types between adjacent and tumor tissues. *P* values were determined by T-test. *, *P* < 0.05. **, *P* < 0.01. ***, *P* < 0.001. **C.** Survival analysis of *IFI30*^+^ macrophages in the E-MTAB-6389 cohort (n = 105). *P* values were determined by T-test. The mean *IFI30*^+^ macrophages score provided by coxph is used to split samples. Samples with score greater than the mean score of all samples are labeled as “*IFI30*^+^ mac high”, and the others are labeled as *IFI30*^+^ mac low”.

**Figure S9.**
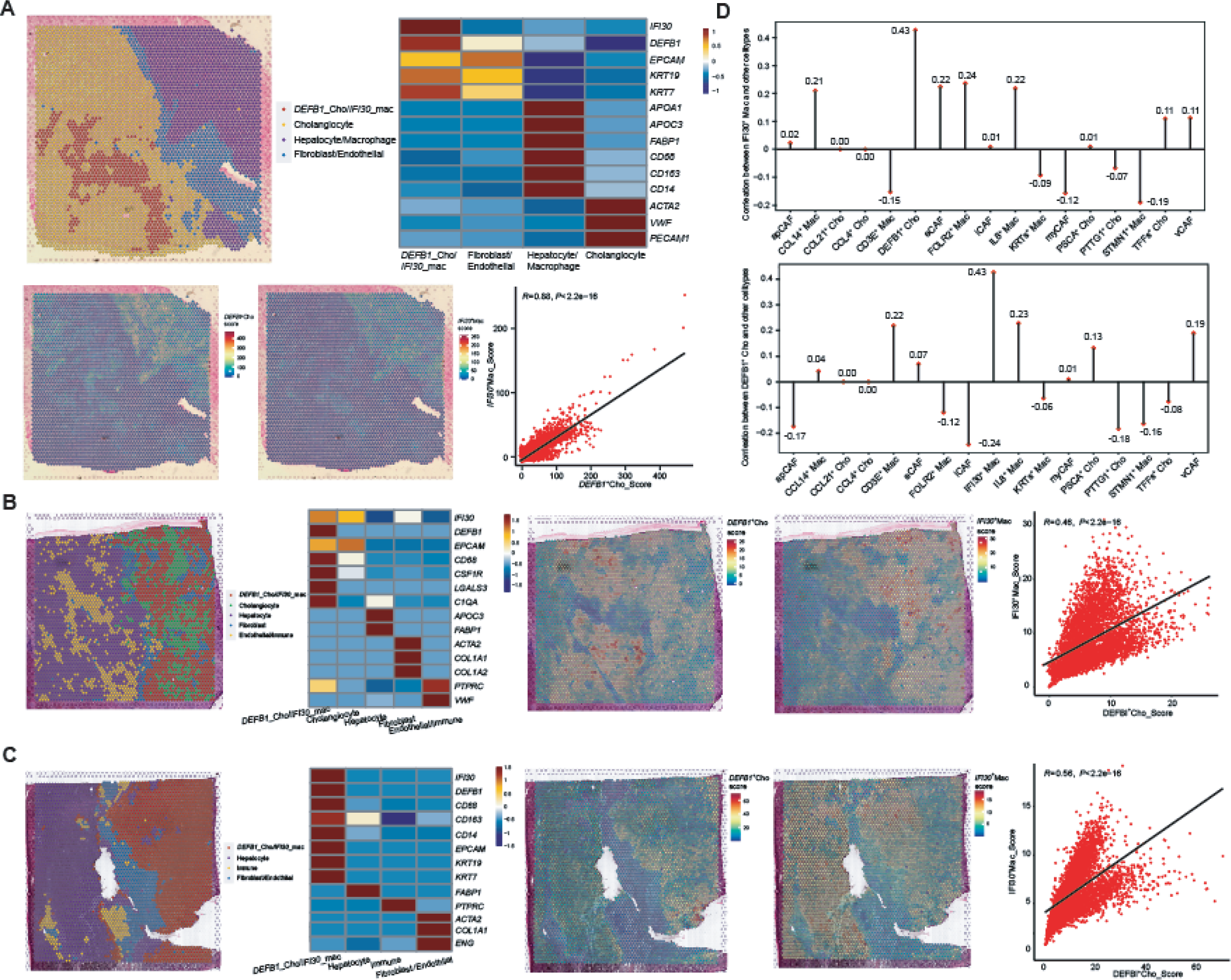
*DEFB1*^+^ cholangiocytes and *IFI30*^+^ macrophages exhibited co-localization in ICC tissue. **A.** Unbiased clustering of spots in spatial sample ICC2, with cell type definition for each cluster. The heatmap shows the expression levels of known markers, spatial feature maps display the feature scores of *DEFB1*^+^ cholangiocytes and *IFI30*^+^ macrophages, and the dot plot shows the correlation of feature scores between the two cell types. *P* values were determined by T-test. **B.** Unbiased clustering of spots in spatial sample ICC3, with cell type definition for each cluster. The heatmap shows the expression levels of known markers, spatial feature maps display the feature scores of *DEFB1*^+^ cholangiocytes and *IFI30*^+^ macrophages, and the dot plot shows the correlation of feature scores between the two cell types. *P* values were determined by T-test. **C.** Unbiased clustering of spots in spatial sample ICC4, with cell type definition for each cluster. The heatmap shows the expression levels of known markers, spatial feature maps display the feature scores of *DEFB1*^+^ cholangiocytes and *IFI30*^+^ macrophages, and the dot plot shows the correlation of feature scores between the two cell types. *P* values were determined by T-test. **D.** The above panel shows infiltration correlation between *IFI30*^+^ Mac and other celltypes. The below panel shows infiltration correlation between *DEFB1*^+^ Cho and other celltypes.

**Figure S10.**
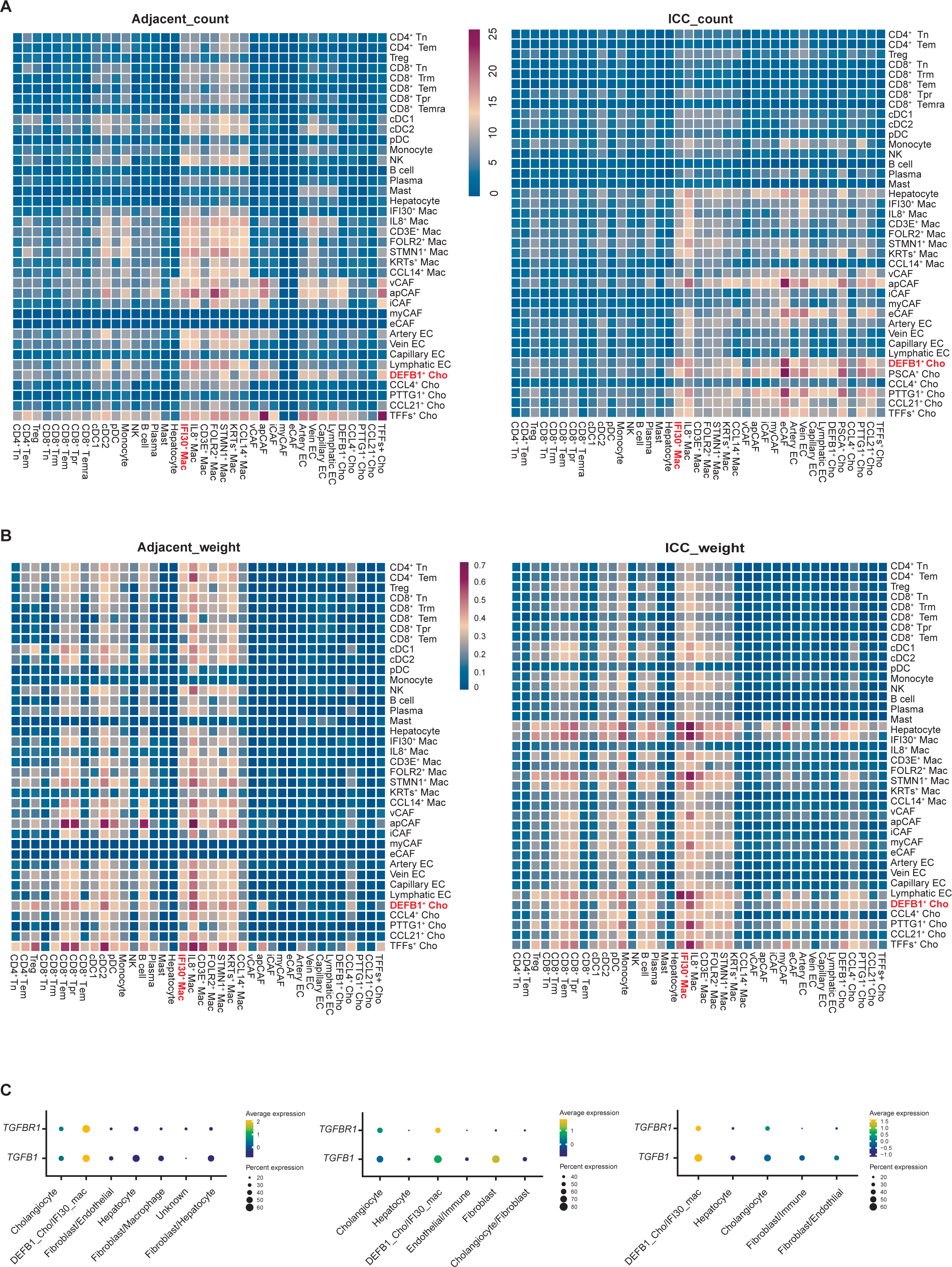
The *TGFB1*/*TGFBR1* pathway is enriched in the interaction between *DEFB1*^+^ cholangiocytes and *IFI30*^+^ macrophages. **A.** Differences in interaction number between cell types in tumor and peritumoral tissues. **B.** Differences in interaction strength between cell types in tumor and peritumoral tissues. **C.** The expression levels of *TGFB1* and *TGFBR1* among cell types in the spatial samples (ICC2, ICC3 and ICC4). The left, middle, and right panels show the result in ICC2, ICC3, and ICC4, respectively.

